# Growth factor-mediated coupling between lineage size and cell fate choice underlies robustness of mammalian development

**DOI:** 10.1101/2019.12.27.889006

**Authors:** Néstor Saiz, Laura Mora-Bitria, Shahadat Rahman, Hannah George, Jeremy P Herder, Jordi García-Ojalvo, Anna-Katerina Hadjantonakis

## Abstract

Precise control and maintenance of the size of cell populations is fundamental for organismal development and homeostasis. The three cell types that comprise the mammalian blastocyst-stage embryo are generated in precise proportions and over a short time, suggesting a size control mechanism ensures a reproducible outcome. Guided by experimental observations, we developed a minimal mathematical model that shows growth factor signaling is sufficient to guarantee this robustness. The model anticipates, without additional parameter fitting, the response of the embryo to perturbations in its lineage composition. We experimentally added lineage-restricted cells to the epiblast both *in vivo* and *in silico*, which resulted in a shift of the fate of progenitors away from the supernumerary cell type, while eliminating cells using laser ablation biased the specification of progenitors towards the targeted cell type. Finally, we show that FGF4 couples cell fate decisions to lineage composition through changes in local concentration of the growth factor. Our results provide a basis for the regulative abilities of the mammalian embryo and reveal how, in a self-organizing system, individual cell fate decisions are coordinated at the population level to robustly generate tissues in the right proportions.

## Introduction

Across metazoa, coordination between cell fate specification and population size ensures robust developmental outcomes. Integration of cell behavior at the population level allows a coordinated response to injury in both embryos and adults (Chen et al., 2015; Wojcinski et al., 2017; Young et al., 2019). The preimplantation mammalian embryo is a paradigm of self-organization, where patterning and morphogenesis occur without the need for maternal determinants or external cues. It therefore provides an *in vivo* platform to understand the processes that ensure precision and robustness during the development of multicellular organisms. These embryos can tolerate cell loss, exemplified by preimplantation genetic diagnose (Harper and SenGupta, 2012), and can incorporate foreign cells to generate chimeric animals (Bradley et al., 1984; Gardner, 1968; Mintz, 1964; Mintz and Illmensee, 1975; Tachibana et al., 2012; Tarkowski, 1959; 1961). Remarkably, neither of these perturbations impair embryonic development. This evidence suggests there are mechanisms that coordinate patterning and population size to enable adaptation. Despite recent interest in understanding and exploiting the capacity of early mammalian embryos and cells for self-organization (Bedzhov and Zernicka-Goetz, 2014; Deglincerti et al., 2016; Harrison et al., 2017; Morgani et al., 2018a; Rivron et al., 2018; Shahbazi et al., 2019; Sozen et al., 2018; Warmflash et al., 2014), little is known about the local control mechanisms that enable such robust autonomous development.

The blastocyst-stage embryo is the hallmark of mammalian preimplantation development. It comprises three cell types – the pluripotent epiblast, which gives rise to the fetus, and the extra-embryonic trophectoderm (TE) and primitive endoderm (PrE, or hypoblast), which predominantly form supporting tissues (Gardner and Rossant, 1979; Kwon et al., 2008; Nowotschin et al., 2019; Papaioannou, 1982; Viotti et al., 2014). In the mouse, these lineages are specified during the 48 hours between embryonic day (E) 2.5 and E4.5, the time of implantation. Epiblast and PrE cells arise from a population of bipotent progenitors and comprise the inner cell mass (ICM) of the blastocyst (Chazaud et al., 2006; Plusa et al., 2008). In the mouse, epiblast specification is driven by the transcription factors NANOG, SOX2 and OCT4 (Avilion et al., 2003; Chambers et al., 2003; Mitsui et al., 2003; Nichols et al., 1998). PrE specification is driven cell-autonomously by GATA6 (Bessonnard et al., 2014; Schrode et al., 2014), which requires activation of the mitogen-activated protein kinase (MAPK) cascade downstream of fibroblast growth factor (FGF) receptors 1 and 2, stimulated by FGF4 (Brewer et al., 2015; Chazaud et al., 2006; Kang et al., 2017; 2013; Krawchuk et al., 2013; Meng et al., 2018; Molotkov et al., 2017; Nichols et al., 2009; Yamanaka et al., 2010).

For development to proceed, the three cell types in the blastocyst must be specified in appropriate numbers and in a short timespan (48h in the mouse, 72-96h in humans). In the mouse embryo, uncommitted ICM progenitors, which co-express NANOG and GATA6, adopt epiblast or PrE identity asynchronously and irreversibly over the course of blastocyst development (Nichols et al., 2009; Saiz et al., 2016b; Schrode et al., 2014; Xenopoulos et al., 2015). In wild type embryos, epiblast and PrE are generated in precise proportions, irrespective of the absolute size of the embryo or the ICM (Saiz et al., 2016b). By contrast, loss of key regulators such as *Nanog*, *Gata6*, *Fgf4* or *Fgfr1* alter these proportions and cause peri-implantation lethality (Bessonnard et al., 2014; Brewer et al., 2015; Frankenberg et al., 2011; Kang et al., 2013; 2017; Krawchuk et al., 2013; Messerschmidt and Kemler, 2010; Mitsui et al., 2003; Molotkov et al., 2017; Schrode et al., 2014; Silva et al., 2009). The ratio of these lineages is likely critical for development of the embryo beyond implantation, and therefore ICM composition must be precisely regulated (Saiz et al., 2016b). However, the details of this tissue size control mechanism remain unclear.

In this study, we combine manipulations of ICM composition with predictions from *in silico* simulations to address this question. We develop a minimal mathematical model in which cell fate decisions in the ICM are mediated solely by intercellular signaling. In this model, ICM cells spontaneously and robustly segregate into two lineages, which scale with embryo size as *in vivo*. The model has only two free parameters, which are adjusted to recapitulate the observed wild type behavior. The robustness of this *in silico* decision is evidenced by the response of the system to perturbations that alter lineage composition. Specifically, the model predicts (with no additional parameter fitting) that reducing or increasing the number of cells in one lineage, would change the pattern of progenitor differentiation to restore lineage composition. This effect is also observed experimentally by using two-photon laser excitation for ablation of specific cells in embryos, and by adding exogenous, lineage-restricted cells to make chimeras. The ability to recover from these perturbations is reduced over time, as the number of uncommitted progenitors is depleted. Finally, we alter the size of the PrE by experimentally tuning the size of the epiblast compartment. Using this system, we show that FGF4 is the growth factor providing the feedback necessary to couple lineage size with cell fate decisions. Our results provide a mechanistic basis for the regulative and scaling abilities of the early mouse embryo and illustrate how a self-organizing system can develop robustly and reproducibly without the need for external inputs.

## Results

### Cell fate decisions in the inner cell mass of the blastocyst are made at the population level

Epiblast and PrE cells originate from a population of bipotent progenitor cells that co-express the lineage-associated transcription factors NANOG (epiblast) and GATA6 (PrE) (Chazaud et al., 2006; Plusa et al., 2008; Saiz et al., 2016b), which we refer to as double positive (DP) cells (Fig. S1A-C). These markers can be used to automatically classify blastocyst cell types and quantify population size (Saiz et al., 2016b; 2016a). PrE cells identified this way express later markers, such as SOX17 and GATA4, in a pattern consistent with previous observations (Fig. S1C-E) (Artus et al., 2011; Kurimoto et al., 2006; Niakan et al., 2010; Nowotschin et al., 2019; Plusa et al., 2008).

Epiblast and PrE size scale with embryo size to maintain a consistent ICM composition (Saiz et al., 2016b). To determine whether this scaling is the result of an active control mechanism or a probabilistic process, we designed a biological probability test in which we mixed labelled (GFP+) with unlabeled cells (GFP-) from 8-cell stage embryos to generate series of chimeric embryos (Fig. 1A; S2A). Using this system, we can alter the fate of one of the two populations and assess the behavior of the other one. If the epiblast and PrE lineage decisions are independent events (i.e., cell-autonomous), the differentiation pattern of the progeny of either cell population (GFP+ or GFP−) should be unaffected by the pattern of the other one. By contrast, if they are not, the probability of any cell adopting a particular fate will be conditional on the fate choice of the others (i.e., a non-cell autonomous decision).

**Figure 1.**
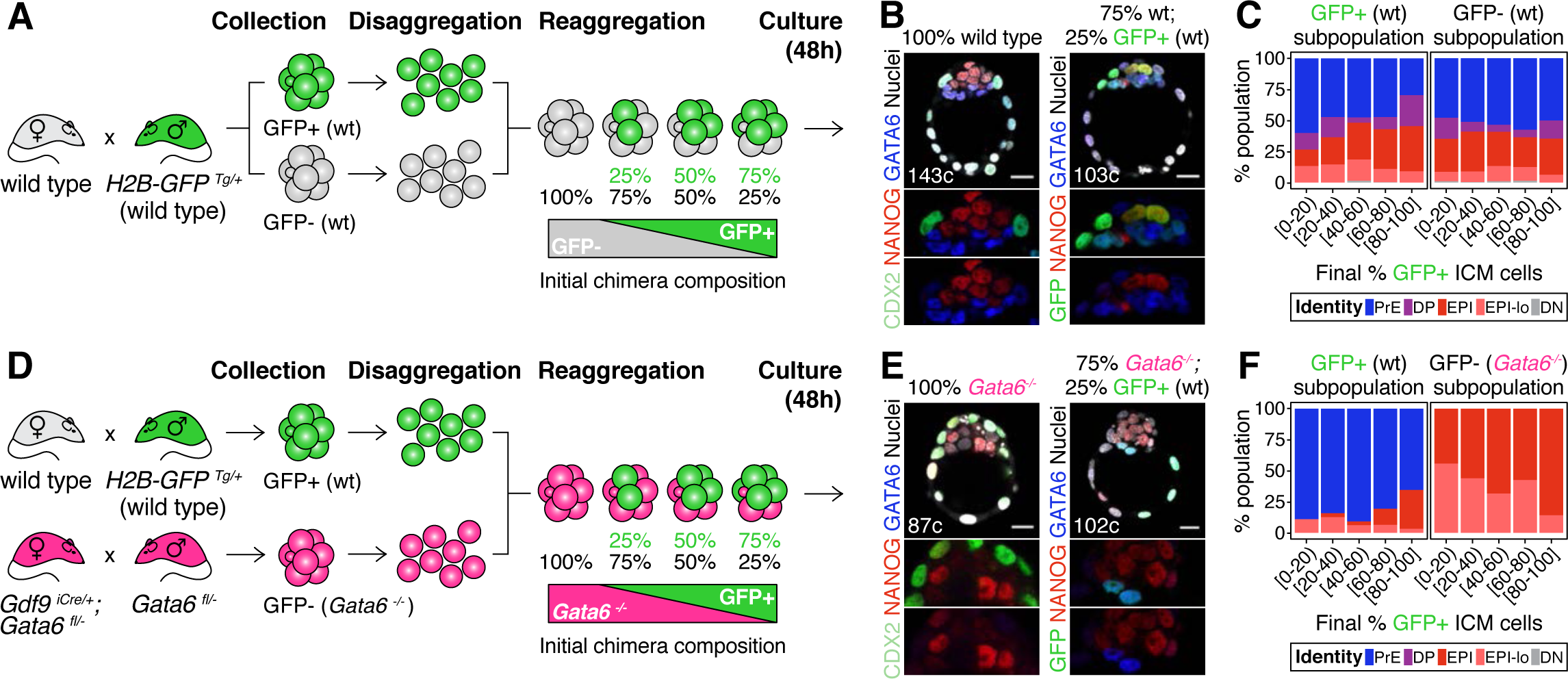
Cell fate decisions in the ICM of the blastocyst are made at the population level. **(A)** Experimental design to test independence of cell fate decisions in the ICM. Wild type embryos (H2B-GFP+ or GFP−) were collected at the 8-cell stage, disaggregated into single cells or clumps, re-aggregated in the combinations indicated and allowed to develop in culture for 48h, until the late blastocyst stage (equivalent to ∼4.5 days post-fertilization). For further experimental details and information on alleles, see Methods. **(B)** Optical cross sections through representative immunofluorescence images of a non-chimeric, wild type control (100% wt) and a chimera made with 75% GFP−; 25% GFP+ cells (all wild type). Magnifications of the ICM are shown below for each marker. **(C)** Stacked bar plots showing the lineage distribution of GFP+ (left) and GFP− (right) wild type ICM cells in embryos, stratified by the final % of H2B-GFP+ ICM cells, as indicated on the x-axis. **(D)** Experimental design to test independence of cell fate decisions in the ICM. Wild type embryos (H2B-GFP+) and *Gata6^−/−^* embryos were collected at the 8-cell stage, disaggregated into single cells or clumps, re-aggregated in the combinations indicated and allowed to develop in culture for 48h, until the late blastocyst stage (equivalent to ∼4.5 days post-fertilization). For further experimental details and information on alleles, see Methods. **(E)** Optical cross sections through representative immunofluorescence images of a non-chimeric, *Gata6^−/−^* control (100% *Gata6^−/−^*) and a chimera made with 75% *Gata6^−/−^*; 25% GFP+ (wt) cells. Magnifications of the ICM are shown below for each marker. **(F)** Stacked bar plots showing the lineage distribution of GFP+ (wt, left) and GFP-*Gata6^−/−^* (right) ICM cells in embryos, stratified by the final % of H2B-GFP+ ICM cells, as indicated on the x-axis. Color coding is indicated. All embryos labeled for NANOG (red), GATA6 (blue) and either CDX2 (controls) or GFP (chimeras) (green). All optical cross sections are 5µm maximum intensity projections. Total cell counts are indicated for each embryo within the merged images. PrE: Primitive Endoderm, DP: Double Positive (for NANOG and GATA6), EPI: Epiblast, EPI-lo: low NANOG epiblast, DN: Double Negative (for NANOG and GATA6). Scale bars = 20µm.

We first mixed labelled and unlabeled wild type (wt) cells, which have unrestricted differentiation potential, in different proportions (Fig. 1A; S2A). The progeny of each population contributed proportionally to TE and ICM (Fig. S2B, C) and to all ICM cell types (Fig. 1B,C; Fig. S2D-F), irrespective of their representation in the resulting chimera. Chimeras made by aggregating two intact 8-cell stage embryos (2x size) or two half embryos (1x size) showed equivalent distributions of cells (Fig. S2A-E), further indicating that lineage allocation is independent of absolute embryo size. Next, we fixed the probability of differentiation of the GFP-population by using *Gata6^−/−^* cells, instead of wt cells (Fig. 1D), and monitored the differentiation pattern of the GFP+ population (wt). *Gata6^−/−^* embryos are cell-autonomously unable to specify PrE cells, and exclusively form TE and epiblast (Fig. 1E, left) (Bessonnard et al., 2014; Schrode et al., 2014). When combined with wt GFP+ cells (Fig. 1D), *Gata6^−/−^* cells give rise to morphologically normal chimeras (Fig. 1E, right; S2G-I). In these chimeras, *Gata6^−/−^* ICM cells make only epiblast, as expected (Fig. 1F, right; Fig. S2J, K), however, the differentiation pattern of the wt compartment (GFP+) changes: wt cells now become biased towards PrE (Fig. 1F, left; Fig. S2K), despite having unrestricted differentiation potential. Moreover, the fate of the wt ICM cells depends on the number of *Gata6^−/−^* cells present: in chimeras with 40% or more mutant cells (< 60% wt GFP+ cells), wt cells contribute almost exclusively to the PrE, whereas in chimeras with fewer mutant cells (> 60% wt GFP+ cells), wt cells contribute to both the epiblast and PrE, and generate ICMs with a normal composition (Fig. 1F, left; Fig. S2K, L).

Taken together, these results show that the chance of wt ICM cells adopting epiblast or PrE fate is conditional on the fate choice made by other ICM cells (Fig. S2M,N). Thus, lineage decisions in the ICM are not independent events, but rather are made at the population level, supporting the notion that intercellular communication coordinates cell behavior to ensure an appropriate ICM composition.

### A minimal model of cell fate decisions solely mediated by growth factor signaling explains robust lineage specification in the ICM

The experimental observations described show that cell fate choice in the ICM is non-cell autonomous. Furthermore, the lineage distribution rapidly trends to a well-defined balanced ratio (close to 50:50) of epiblast:PrE cells (Fig. S1A, B; S3A-C), characterized by embryo-to-embryo variability that decreases over time (Fig. S3D). From a dynamical systems perspective, these features are indicative of the existence of a balanced attractor state within the population of proliferating cells. To test whether cell-cell signaling is sufficient to generate such an attractor, we developed a mathematical model (see Box 1) in which a cell fate switch depends on the levels of a growth factor, which is under the explicit control of a lineage-specific transcription factor. In the context of the blastocyst, FGF4 and NANOG, respectively, fit these categories (Fig. 2A and Supplementary Text), although this model can be generalizable to any circuit with similar characteristics.

**Figure 2.**
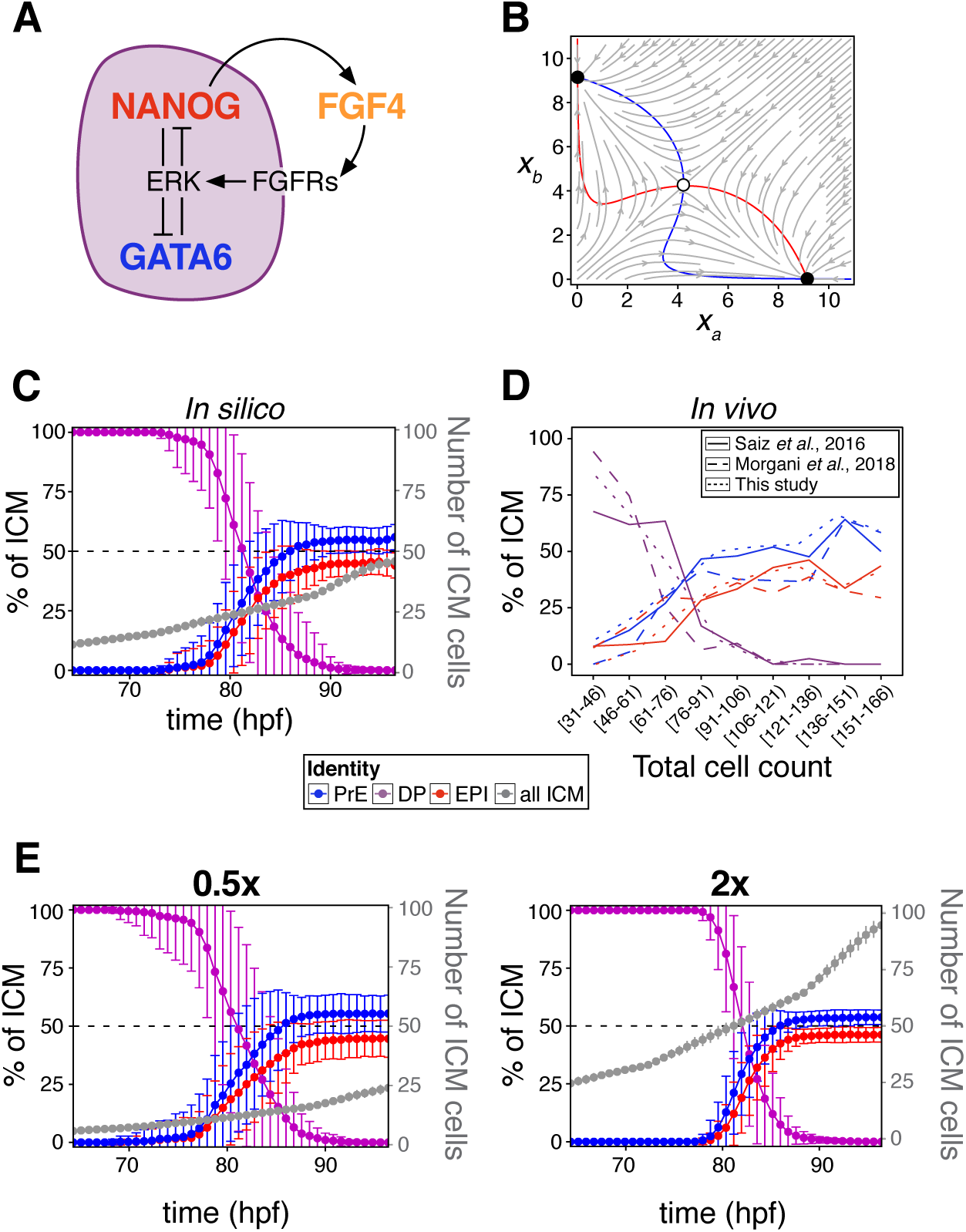
A minimal model of cell fate decisions solely mediated by growth factor signaling explains robust lineage specification in the ICM. **(A)** Diagram of our proposed model of molecular control of cell fate in the ICM. **(B)** Phase plane of our model for the case of a two-cluster state. Each axis shows the NANOG levels in one of the two clusters. Red and blue lines show the nulclines of the system (Box 1); the gray arrows depict typical trajectories of the system; and black (white) circles correspond to stable (unstable) equilibria of the two-cluster system. **(C)** Lineage dynamics in *in silico* simulations of ICM development using our proposed model. (D) *In vivo* ICM lineage dynamics from three experimental datasets, as indicated. **(E)** Lineage dynamics in *in silico* simulations of scaling experiments (to be compared with the experimental results of (Saiz et al., 2016b)). Absolute ICM size was modified to 0.5 or 2x the normal size, as indicated.

In our model, the state of the system is described by a single variable *x_i_* per cell, which in our case can be considered to represent the amount of NANOG in that cell. FGF4 is assumed to be activated by NANOG and feeds back onto NANOG and GATA6 via ERK (see Supplementary Text). As a result, cell-cell signaling effectively drives an indirect mutual inhibition between NANOG and GATA6, in such a way that the level of NANOG in a given cell is inhibited by that in neighboring cells (see Box 1), a mechanism reminiscent of lateral inhibition. As in lateral inhibition, the model exhibits an attractor state in which two cell clusters coexist (Collier et al., 1996), corresponding to states of low and high NANOG (high and low GATA6, respectively). The simplicity of this system allows us to interpret its behavior geometrically, in analogy with earlier work on the interaction between EGF and Notch signaling (Corson and Siggia, 2017). The associated phase-plane portrait in our case is shown in Fig. 2B, in which the two stable states are represented by black circles towards which NANOG levels tend with time (gray arrows). The situation depicted in Fig. 2B corresponds to a perfectly symmetric case in which the epiblast:PrE ratio is 50:50 and cells are perfectly mixed (see Box 1), and as we go on to show the attractor is robust to variations in this perfect balance.

We next sought to determine how cell fate decisions are established according to the model, when the cells proliferate and are rearranged as the embryo develops and grows. To that end, we implemented an agent-based model to simulate the growth of the ICM (see Supplementary Text), in which cells divide and interact with one another via a soft-sphere potential (Tosenberger et al., 2017). The biochemical model described is applied to the interacting cells, and the system is allowed to progress biochemically and dynamically until a fixed cell number is reached. Simulations show that a stable cell-type distribution is consistently reached (Fig. 2C; Movie S1), in agreement with the described attractor dynamics, and in line with experimental observations from this study and others (Fig. 2D). The decision is robust, since altering the absolute cell number of this *in silico* ICM leads to scaling of lineage size to maintain ICM composition (Fig. 2E), in agreement with experimental observations (Saiz et al., 2016b). These results suggest that growth factor-mediated feedback is sufficient to endow the embryo with robustness to perturbations. Reaching this conclusion required us to uncouple cell-cell signaling interactions from intracellular (cell-autonomous) mechanisms, something not possible experimentally, *in vivo*, and which would be challenging to do in more detailed biochemical models (Bessonnard et al., 2014; De Mot et al., 2016; Tosenberger et al., 2017).

#### Box 1. A minimal model of growth factor-mediated cell-fate decisions

We use the following model to represent the dynamics of a population of cells in which the transcription factors NANOG and GATA6 mutually repress each other through extracellular growth factor signaling:

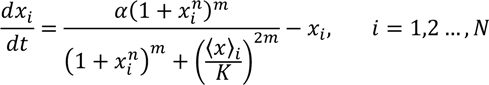

Here *x_i_* represents the (dimensionless) concentration of NANOG in cell *i*, and ⟨*x*⟩*_i_* denotes the average value of *x* over the immediate neighborhood of cell *i*, including *x_i_* itself. The dynamics described by the equation above corresponds to motion in a potential that is double-well shaped for large enough values of the mean field ⟨*x*⟩, which is the situation of the DP fate. The two potential wells represent the high and low NANOG states, and therefore, our model implies that the DP fate is not a stable equilibrium of the cell, but a transient state towards a stable epiblast or PrE fate.

This model can be derived from a specific molecular-level circuit explicitly involving NANOG, GATA6 and the growth factor FGF4, and implicitly ERK signaling downstream of FGF receptors, as detailed in the Supplementary Text. However, due to its minimal character, the model is not unique to the molecular interactions assumed to be involved in this cell fate decision; other molecular circuits can likely be reduced to it to achieve the same result.

In the model, the level of NANOG is inhibited by that of its neighbors, in a manner that resembles lateral inhibition-mediated signaling. We ask whether such a model can sustain solutions in which cells cluster into two distinct cell types, expressing NANOG at two different levels (which we could interpret as epiblast and PrE cells). Epiblast and PrE cells display a salt-and-pepper distribution in the ICM at early blastocyst stages (Chazaud et al., 2006). Therefore, we assume perfect mixing among the cells within the population, in which case such a two-cluster state would be described by the following two-dimensional dynamical system:

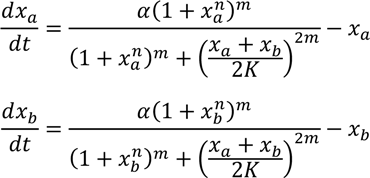

The dynamics of this potential two-cluster solution can be examined via the phase-plane portrait shown in Fig. 2B, which displays the nullclines of the system in which each of the derivatives is zero (in red and blue), the typical flow exhibited by the system from different initial conditions (in gray) and the equilibrium points of the system, two of them stable (black circles) and the third one unstable (white circle). Note that the two stable equilibria correspond to the same behavior of the system, since the labels *a* and *b* of the two clusters are interchangeable.

The existence of the two symmetric stable equilibria ensures that the two-cluster state is a solution of the system, and that the population splits spontaneously into two distinct fates, as we show by means of agent-based simulations (described in the Supplementary Text) throughout the text.

### The lineage composition of the ICM is robust to expansion of the epiblast

The attractor solution found in the model described suggests that the cell-fate decision reached by the embryo is robust to perturbations, in particular to those affecting the size of the population. To probe the capacity of the system to perceive and adjust for changes in cell numbers, we expanded the epiblast by introducing increasing amounts of mouse embryonic stem cells (ESCs) into embryos (Fig. 3A). ESCs are derived from the epiblast and contribute to this lineage when re-introduced into an embryo (Beddington and Robertson, 1989; Boroviak et al., 2014; Brook and Gardner, 1997; Lallemand and Brûlet, 1990; Nagy et al., 1990; Tokunaga and Tsunoda, 1992). To control for variation in the contribution of ESCs to chimeras, we stratified embryos based on the final size of the ESC compartment, relative to the average size of the epiblast in controls (1x control EPI = 5-10 cells, 2xEPI = 10-20 cells, etc.) (Fig. S4A). This way we generated chimeras with increasingly larger epiblasts and asked how ICM composition was affected (Fig. 3B; S4B; Movies S2, S3).

**Figure 3.**
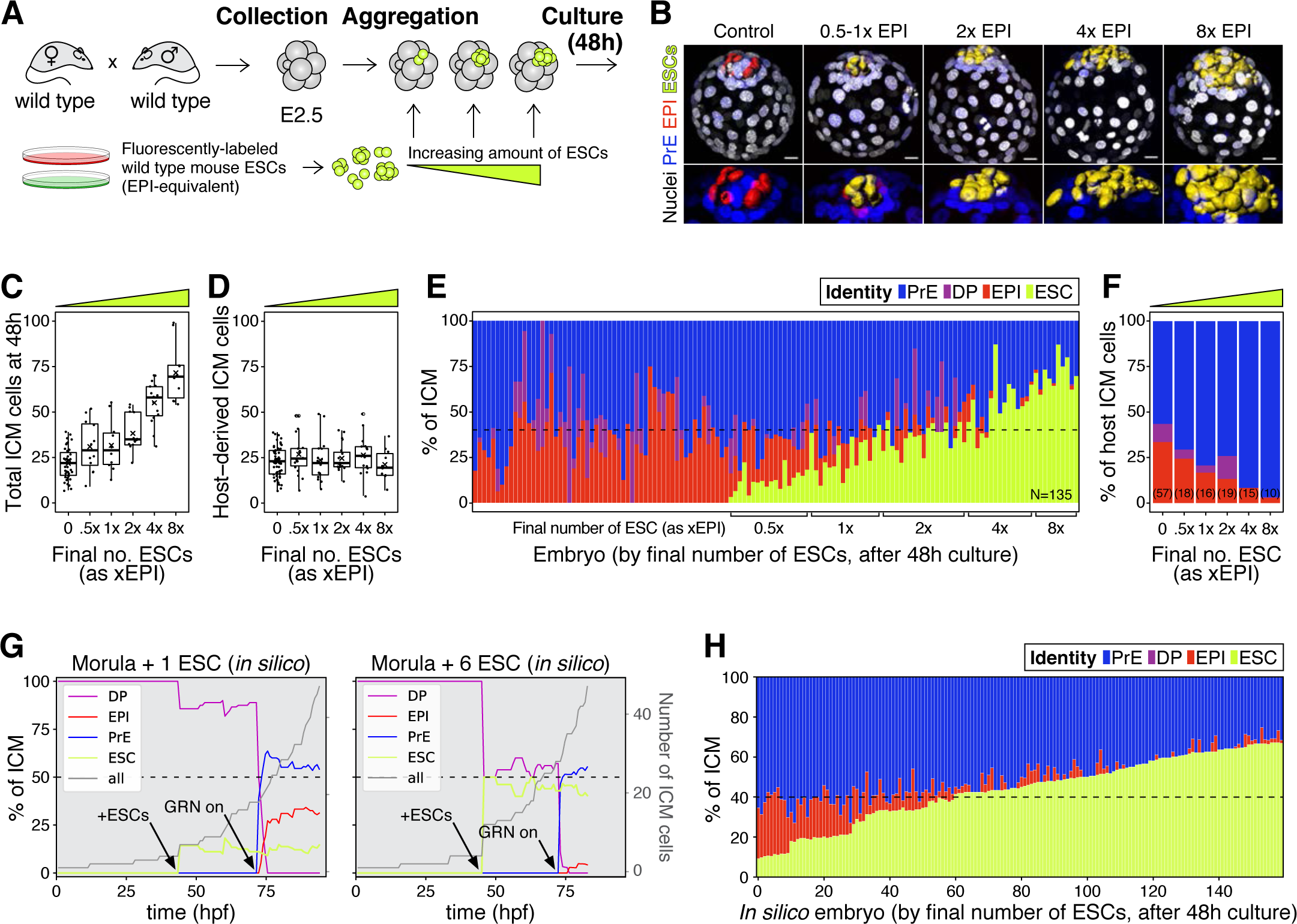
The lineage composition of the ICM is robust to expansion of the epiblast. **(A)** Experimental design. 8- or 8-16 cell stage embryos (2.5 days post-fertilization) were recovered from CD1 (wild type) females crossed with CD1 males. Embryos were denuded, aggregated with clumps of fluorescently labelled ESCs and cultured for 48-56h, until the late blastocyst stage (equivalent to ∼4.5 days post-fertilization). **(B)** 3D renders of a series of control (no ESCs) and chimeras, generated as indicated in (A), carrying increasing amounts of ESCs (indicated as EPI-equivalent size (x EPI)). Control epiblast and ESCs in chimeras are highlighted as computer-rendered volumes and color coded as indicated. GATA6+ PrE is shown in blue, NANOG+ host epiblast is shown in red where applicable. **(C)** Box plot showing the total size of the ICM (host-derived + ESCs) at the end of the experiment in each group of embryos, as defined by the size of the ESC compartment. **(D)** Box plot showing the size of the host-derived ICM component at the end of the experiment in each group of embryos, as in (C). **(E)** Stacked bar plot showing the relative ICM composition at the end of the experiment for all embryos analyzed. Each bar represents the ICM of one embryo, ordered by increasing absolute number of ESCs at the end of the experiment. Dashed line indicates the normal ratio of 60% PrE:40% epiblast found in intact wild type embryos. Number of embryos analyzed is indicated (N). Brackets on x-axis indicate the number of ESCs in those embryos, relative to the size of the average wt control epiblast (xEPI). **(F)** Stacked bar plot showing the relative contribution of host cells to each of the ICM lineages in each group of embryos. Yellow wedge represents the increasing amount of ESCs in each group. **(G)** Growth curves showing lineage dynamics in *in silico* simulations of the aggregation experiments shown in (A). Left Y-axis and curves for each lineage indicate relative size (as % of ICM). Right Y-axis and gray curves indicate total number of ICM cells (including ESCs). **(H)** Stacked bar plot showing the relative ICM composition at the end of the experiment in *in silico* simulations of the experiments shown in (A). Each bar represents the ICM of one simulated embryo (i.e., a single iteration), and bars are arranged by increasing absolute number of ESCs at the end of the simulation, as in (E). Dashed line indicates the normal ratio of 60% PrE:40% epiblast found in intact wild type embryos. Color coding is indicated for (E, G, H). In all box plots whiskers span 1.5x the inter quartile range (IQR) and open circles represent outliers (values beyond 1.5x IQR). Crosses indicate the arithmetic mean and each dot represents one embryo. Yellow wedges represent the increasing amount of ESCs in each group. PrE: Primitive Endoderm, DP: Double Positive (for NANOG and GATA6), EPI: Epiblast, ESC: embryonic stem cell. Scale bars = 20µm.

Increasing the number of ESCs in chimeras increased the overall size of the ICM (host + ESCs) (Fig. 3C), although it did not affect the contribution of the host embryo to the ICM (Fig. 3D) or the ratio between TE and host ICM (Fig. S4C). This result suggests that ESCs have no net effect on the survival or proliferation of host cells. By contrast, increasing the number of ESCs in chimeras reduced the contribution of host cells to the epiblast (Fig. 3E-F; S4D-E) and increased their contribution to the PrE (Fig. 3F; S4F). This shift resulted in maintenance of the ICM composition in chimeras comprising as many ESCs as 2-4xEPI, but not more (Fig. 3E). The ICM composition of chimeras comprising an equivalent of 4xEPI, was in most cases comparable to that of *Gata6^+/-^* or *Fgf4^+/-^* blastocysts (∼40% PrE, 60% EPI; Fig. 3E; S4D), which develop into fertile adults (Kang et al., 2013; Krawchuk et al., 2013; Schrode et al., 2014), suggesting that an expansion of up to 4x epiblast is compatible with normal development.

We next asked whether this experimental perturbation would be recapitulated by our mathematical model *in silico*. To do so, increasing amounts of epiblast-equivalent cells (ESCs) were added to the system before activation of the molecular circuit (Fig. 3G, arrows). These cells produce FGF and thereby impact NANOG dynamics in their neighbors but, being lineage restricted, are not subject to the same regulatory dynamics as host cells. In agreement with our experimental results, these ESCs effectively contribute to the epiblast compartment to maintain the overall ICM composition for a wide range of perturbations (Fig. 3G, H). In particular, as the number of added ESCs is increased, the number of host cells that acquire an epiblast fate is progressively reduced, until the epiblast compartment is eventually composed only of ESCs. Notably, all these perturbations shift cell fate choice in the ICM toward PrE without net ICM growth, indicating there is no compensatory proliferation in this context and that the regulative capacity of the system is mediated only by changes in cell fate allocation. Together, these experimental and modeling results underscore the robustness of the system and further establish a population-level coordination of cell fate choice.

### The lineage composition of the ICM is robust to *in silico* reduction of lineage size

The chimera experiments described allow us to probe the response of the embryo to perturbations at a fixed time, early in the process, but not as progenitors are depleted over time (Fig. S1A, B). However, an attractor state such as the one described in Fig. 2 should also be robust to perturbations all along the system’s trajectory, as long as they are not exceedingly large. Experimentally it is not possible to expand the epiblast using ESCs at sequential stages of blastocyst development (not shown). Instead, to probe the ability of the system to adjust to perturbations over time, we used our model to modify ICM composition *in silico* to different degrees and at sequential time points. With that goal in mind, we first eliminated 30% of the cells of all three ICM lineages (EPI, PrE and DP) at a time when each represents ∼1/3 of the ICM. This perturbation, which is equivalent to scaling down the absolute size of the ICM by 30%, does not alter the relative composition of the ICM (Fig. 4A). As in our simulations of embryo scaling (Fig. 2E), the relative ICM composition at the end of the simulation was maintained robustly across embryos (Fig. 4B). Eliminating cells in this way (equally across cell types) had no effect on ICM composition irrespective of the developmental stage at which it was carried out or the magnitude of the perturbation (Fig. 4C).

**Figure 4.**
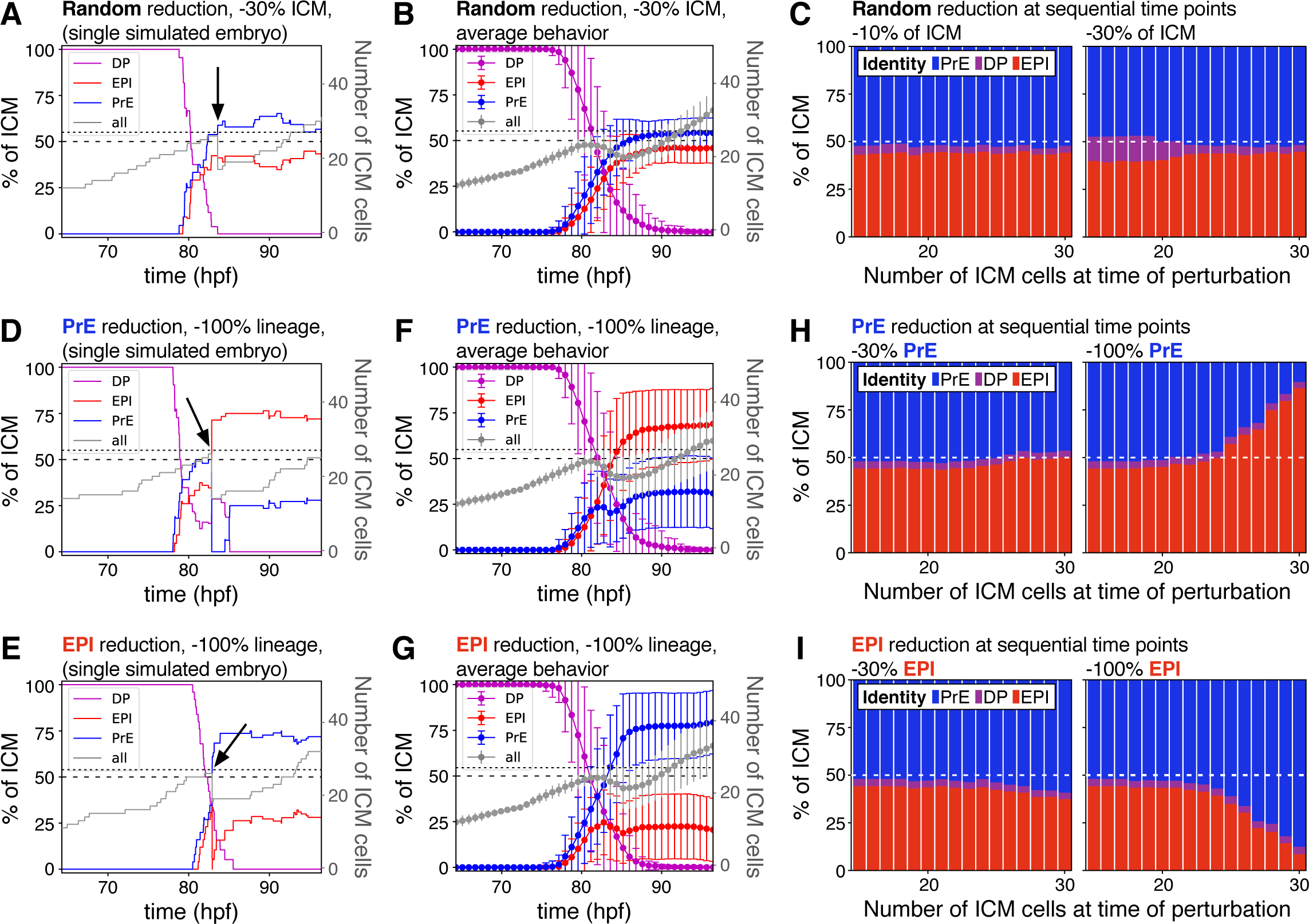
The lineage composition of the ICM is robust to *in silico* reduction of lineage size. **(A)** Growth curves for each ICM lineage after simulation of a 30% reduction in ICM size (3 PrE, 3 DP and 3 epiblast cells removed from a 27-cell ICM) using our model described in Fig. 2. Arrow indicates the time of cell elimination. Lines are color-coded for each lineage, as indicated and represent relative lineage size (scale on the left Y-axis). Grey line indicates the absolute size of the ICM, as shown on the right Y-axis. Dotted line indicates 27 ICM cells. Dashed line indicates 50% of the ICM, for reference. **(B)** Growth curves as those in (A) showing the average behavior for 100 simulations. Error bars indicate the standard deviation. **(C)** Stacked bar plots showing the final ICM composition after simulating the elimination of 10% (left) or 30% (right) of ICM cells at sequential points in embryo development. Developmental stage at the time of cell elimination is indicated on the x-axis as number of ICM cells (15-30 ICM cells, equivalent to ∼50-100 total cells). **(D, E)** Growth curves for each ICM lineage after simulation of a 100% reduction in PrE (D) or epiblast (E), when the ICM reaches 27 cells, as shown in (A) and indicated by the arrow. Lines are color-coded for each lineage, as indicated and represent relative lineage size (scale on the left Y-axis). Grey line indicates the absolute size of the ICM, as shown on the right Y-axis. Dotted line indicates 27 ICM cells. Dashed line indicates 50% of the ICM, for reference. **(F, G)** Growth curves as those in (D, E) showing the average behavior for 100 simulations of PrE (F) and epiblast (H) reduction. Error bars indicate the standard deviation. **(H, I)** Stacked bar plots showing the final ICM composition after simulating the elimination of 30% (left) or 100% (right) of the PrE (H) or the epiblast (I) at sequential points in embryo development. Developmental stage at the time of cell elimination is indicated on the x-axis as number of ICM cells (15-30 ICM cells, equivalent to ∼50-100 total cells). Color coding is indicated. hpf: hours post-fertilization, PrE: Primitive Endoderm, EPI: epiblast, DP: double positive.

We then eliminated PrE or epiblast-biased cells only, therefore causing an acute deviation in the normal ICM ratio. When we removed 100% of either lineage at the same developmental stage as above, we observed only a partial recovery of the targeted lineage (Fig. 4D-G). However, a partial reduction (−30%) in either lineage was completely compensated for, and normal ICM composition was restored at all developmental stages (Fig. 4H-I, left). By contrast, the ability to recover from loss of 100% of a lineage was reduced over time, as progenitor cells are lost (Fig. 4H-I, right), indicating that the ability of the system to recover from changes in ICM composition depends on both the presence of uncommitted progenitor cells and the magnitude of the perturbation.

### Laser ablation enables alteration of ICM composition with high spatiotemporal control in mouse embryos

To experimentally validate the predictions of our mathematical model, we used a multi-photon laser to eliminate cells and thus alter lineage composition in live embryos (Fig. 5A, C and see Methods). Laser cell ablation is routinely used in non-mammalian systems to eliminate cells in a non-invasive way. In mammalian embryos, however, while it has been previously applied to disrupt tissues (Eiraku et al., 2011; Reupke et al., 2009; Takaoka et al., 2017), its potential to target single cells and its effect on developmental competence has not been determined. We combined two spectrally distinct nuclear reporters to identify all cell types in the ICM: a reporter for *Pdgfra* expression (*Pdgfra^H2B-GFP/+^* (Hamilton et al., 2003; Plusa et al., 2008)) and a ubiquitous nuclear mKate2 reporter (Susaki et al., 2014). PrE and uncommitted progenitors were labelled with different levels of GFP (Plusa et al., 2008; Xenopoulos et al., 2015), whereas epiblast cells are identified by the presence of mKate and the absence of GFP (Fig. 5B, D; S5A-D; Movies S4-5) (see Methods for details on cell identification and classification). We verified the death of targeted cells using time-lapse imaging (Fig. 5E; Movie S6) and estimated their half-life to be 3.4h (Fig. 5F). The use of a two-photon laser allowed targeting of individual nuclei on any z-plane of the ICM without affecting other cells in the light path. Untargeted cells in both ablated and control embryos showed identical survival (Fig. 5F), including those immediately adjacent to targeted cells (Fig. 5G). The gross morphology and size of ablated embryos was comparable to that of intact controls, even after targeting 100% of either lineage (Fig. 5H; S6A).

**Figure 5.**
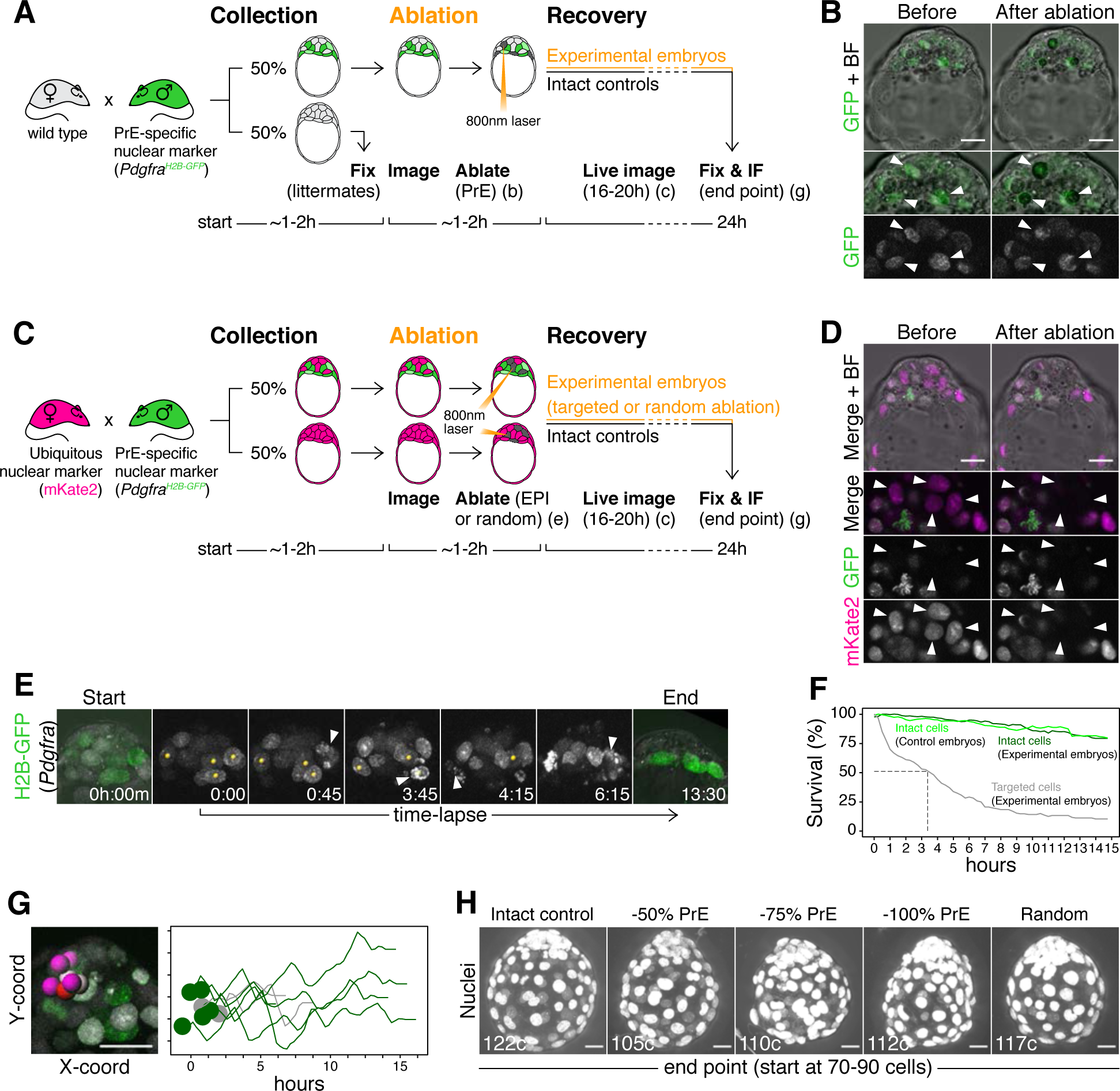
Laser ablation enables alteration of lineage size with high spatiotemporal control in mouse embryos. **(A)** Experimental design for PrE ablation. Blastocysts recovered from CD1 (wild type) females crossed with *Pdgfra^H2B-GFP/+^* or *Pdgfra^H2B-GFP/+^; R26:CAG:3x-nls-mKate2^Tg/Tg^* males were sorted for GFP. GFP-embryos were fixed as reference littermates to estimate the developmental stage of the entire litter. GFP+ embryos were used for the experiment and subject to ablation of different amounts of PrE cells (GFP+) followed by 16-20h of live imaging to visualize the response to cell ablation, followed by 8-4h of *in vitro* culture (for a total of 24h). **(A)** Representative images of live GFP+ embryos before and after ablation of PrE cells. Bottom panels show magnifications of the ICM with GFP alone on grayscale, as indicated. Arrowheads point at targeted PrE cells. **(C)** Experimental design for epiblast ablation. Blastocysts were recovered from *R26:CAG:3x-nls-mKate2^Tg/Tg^* females crossed with *Pdgfra^H2B-GFP/+^; R26:CAG:3x-nls-mKate2^Tg/Tg^* males. GFP-, mKate2+ embryos were used as either intact or random ablation controls. GFP+, mKate2+ embryos were used for the experiment and subject to ablation of different amounts of epiblast cells (GFP-) or PrE cells (GFP+), followed by 16-20h of live imaging to visualize the response to cell ablation, followed by 8-4h of *in vitro* culture (for a total of 24h). **(D)** Representative images of live GFP+ embryos before and after ablation of epiblast cells. Bottom panels show magnifications of the ICM, for both markers together and each of them on grayscale, as indicated. Arrowheads point at targeted epiblast cells. **(E)** Still images of a representative embryo in the hours after ablation. See also Movie S4. Yellow spots on grayscale images mark targeted cells. Arrowheads point at cell death of each targeted cell. All images are timestamped as h:mm. **(F)** Survival curves for targeted (gray) and intact cells in both experimental (dark green) and intact control embryos (light green). Dashed line marks half-life of targeted cells (∼3h) **(G)** Survival of intact cells neighboring targeted cells. Image shows selected ICM cells (intact DP and epiblast cells, color coded, and targeted PrE cells, gray). Graph shows X-Y coordinates of cells shown in picture and survival (hours) for each cell. Time scale for each cell is shifted based on their initial X position, for visualization purposes. **(H)** Maximum intensity projections of representative embryos fixed after 24h in culture, showing all nuclei over bright field image. Treatment was done at the 70-90 cell stage and is indicated above. Total cell count for each embryo is shown within each image. PrE: Primitive Endoderm, EPI: Epiblast. Scale bars = 20µm.

### The cell fate choice of uncommitted ICM progenitors is dictated by lineage size

We used laser ablation (Fig. 5A, C) to test how recovery from changes in ICM composition correlates with developmental time and magnitude of perturbation (Fig. 4F-G). We eliminated increasing amounts of PrE or epiblast cells (2-18 cells, representing 50-100% of either lineage) at sequential stages of blastocyst development (embryos comprising 50-110 total cells) and assessed the composition of the ICM after a recovery period of 24h (Fig. 6A). Although embryos showed a reduction in ICM size as a result of the perturbation (Fig. S6B), they contained both PrE and epiblast cells (Fig. 6B), indicating at least a partial ability to recover. Predictably, their ability to re-establish a normal PrE:epiblast ratio was reduced the later the developmental stage at the time of cell ablation (Fig. 6C-D; S6C). As in our simulations (Fig. 4H), this deviation was more pronounced the larger the magnitude of the perturbation (Fig. 6C-D; S6C).

**Figure 6.**
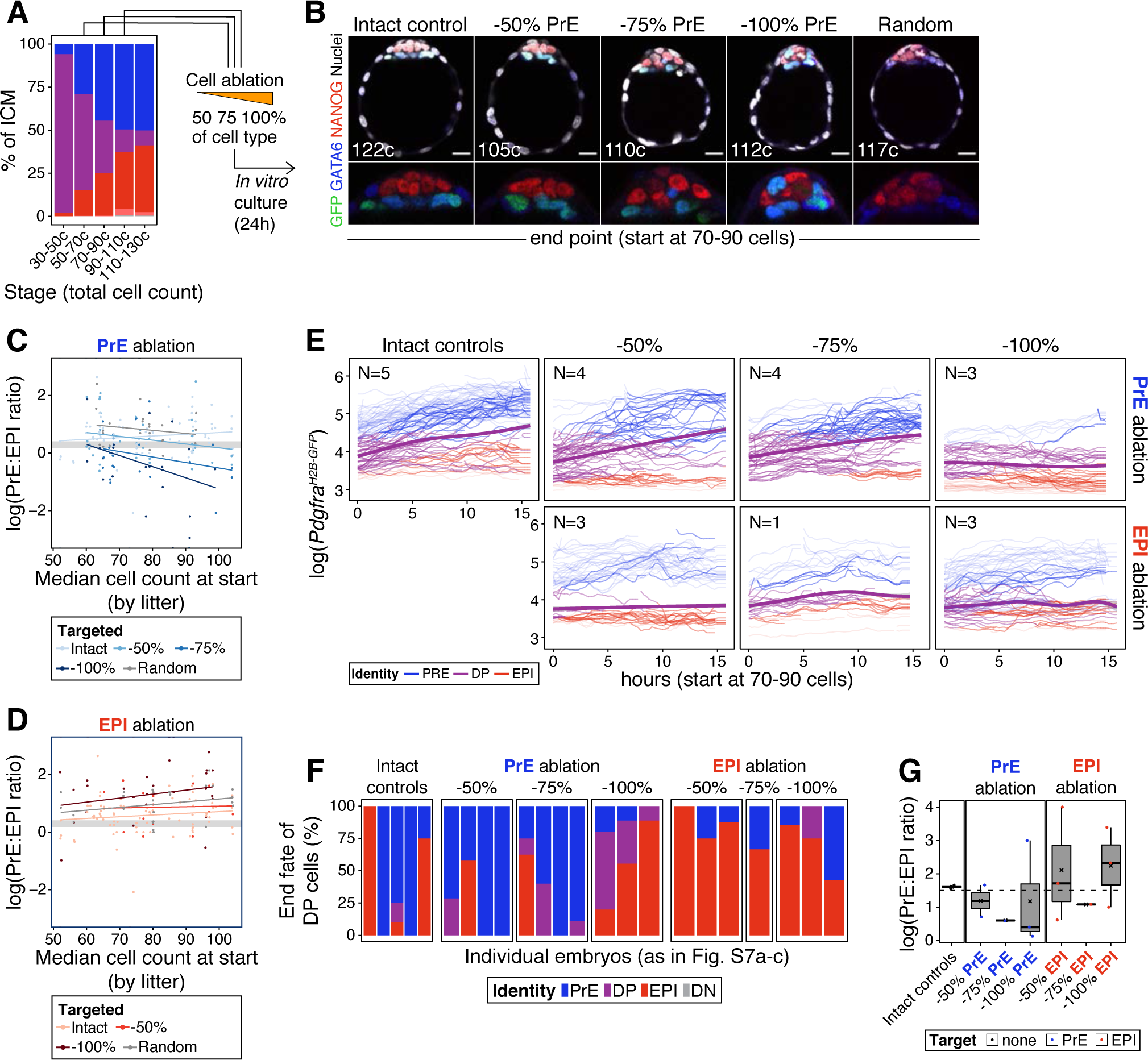
The cell fate choice of uncommitted ICM progenitors is dictated by lineage size. **(A)** Stacked bar plot showing ICM composition at sequential stages of blastocyst development. Embryos at each of these stages were subject to laser ablation of different fractions of either the PrE or the epiblast and allowed to recover *in vitro* for 24h (see Fig. 5A, C). **(B)** Optical cross sections through representative immunofluorescence images of embryos subject to ablation at the 70-90 cell stage and fixed at the end of the experiment (24h later) (same embryos as in Fig. 5G). Embryos are labelled for NANOG (red) and GATA6 (blue) to identify all ICM cell types. H2B-GFP, indicating *Pdgfra* expression, is shown in green where applicable. Lower panels show magnifications of the ICM Treatment for each embryo is indicated over each image. **(C)** PrE:epiblast ratio (shown as natural logarithm) at the end of the experiment (24h) in embryos where fractions of the PrE were eliminated at sequential stages of blastocyst development, as indicated on the x-axis. Shades of blue indicate the magnitude of the reduction in the PrE. Intact controls are embryos in which no cells were killed, Random controls are embryos in which randomly chosen ICM cells were killed without knowing their identity, in equivalent numbers to the −100% group. **(D)** PrE:epiblast ratio (shown as logarithm) at the end of the experiment (24h) in embryos where fractions of the epiblast were eliminated at sequential stages of blastocyst development, as indicated on the x-axis. Shades of red indicate the magnitude of the reduction in the epiblast. **(E-F)** *In silico* simulation of the same experiments shown in (C-D), using our model, described in Fig. 2. **(G)** Dynamics of *Pdgfra^H2B-GFP^* expression in progenitor cells (DP) of experimental embryos targeted at the 70-90 cell stage. Each line represents one cell. Color coding indicates cell identity, as inferred from reporter expression (see Methods). Number of embryos per plot is indicated. Cells classified as PrE or epiblast at the beginning of the experiment are shown as color-coded semi-transparent lines behind progenitor cells, for reference. Smoothing curves for *Pdgfra* expression in progenitor cells are shown as thick purple lines. Fraction of the PrE or epiblast eliminated is indicated above each panel, lineage targeted is indicated to the right of each panel. **(H)** Stacked bar plots showing the final identity adopted by progenitor (DP) cells in each of the embryos plotted in (G). **(I)** Box plots showing the PrE:epiblast ratio (shown as logarithm) at the end of the movie, in embryos where all or most of the ICM cells could be tracked after cell ablation at the 70-90 cell stage (subset of embryos shown in (G) and (H)). Compare to panels (C) and (D). Treatment is indicated on the x-axis. In box plots whiskers span 1.5x the inter quartile range (IQR) and open circles represent outliers (values beyond 1.5x IQR). Cross indicates the arithmetic mean and each dot represents one embryo. PrE: Primitive Endoderm, DP: Double Positive (for NANOG and GATA6), EPI: Epiblast. Scale bars = 20µm.

We next assessed the response of progenitor cells to changes in ICM composition. We tracked individual cells in time-lapse movies for 15h following cell ablation. Progenitor cells in intact embryos contribute to both epiblast and PrE, as assessed by *Pdgfra* expression (Fig. S5F; 6E). We found a bias of these cells towards PrE in intact embryos (Fig. 6E), likely due to oversampling of PrE (given their co-expression of *Pdgfra* and mKate2), as well the overrepresentation of PrE cells within the ICM. Progenitor cells in embryos where the PrE was targeted showed a comparable trend to control embryos, generally upregulating *Pdgfra* expression and acquiring a PrE identity (Fig. 6E, purple line). Conversely, in embryos where epiblast cells were eliminated, progenitors had a propensity to downregulate or maintain low levels of *Pdgfra* and become epiblast (Fig. 6E, purple line). Accordingly, at the end of movies, cells initially identified as progenitors were more frequently found in the lineage that had been targeted (Fig. 6F), and the final PrE:epiblast ratio (in movies where we could track all or most of the ICM cells) was comparable to that obtained with end-point analysis experiments (Fig. 6G and C-D; Fig. S6C and D).

Finally, we investigated the relative contribution of each cell behavior (cell death, division and specification of progenitors) to recovery of ICM composition after cell ablation. In addition to changes in progenitor specification (Fig. 6E, F), we observed both cell proliferation and cell death in the PrE and epiblast compartments for each embryo we could track (Fig. S7A-E). Cell ablation increased the survival of progenitor cells, although it had no notable effect on either intact PrE or epiblast cells (Fig. S7F). Conversely, we observed a slight increase in progenitor proliferation following PrE, but not epiblast ablation (Fig. S7G). Lastly, in the sampled embryos where we targeted 100% of the PrE (N=3), we found a high degree of cell death that was not compensated for by proliferation or progenitor specification (Fig. S7B), and which resulted in an inability to restore ICM composition (Fig. 6E-F).

Overall, these results indicate that changes in ICM composition are primarily compensated for by a shift in the differentiation pattern of progenitor cells. Although the small sample size precludes any definitive conclusion regarding the roles of cell death and proliferation, both our cell ablation and chimera data suggest they play only accessory roles in this process and that uncommitted progenitors are the primary substrate for regulation.

### FGF4 provides the dynamic readout of lineage size that determines cell fate

We have shown here that the lineage composition of the ICM is robust to changes in absolute tissue size, both *in vivo* and *in silico* and propose that growth factor-mediated feedback ensures this robustness (Fig. 2C). Fibroblast growth factor 4 (FGF4), secreted by ICM cells, is necessary for PrE specification (Kang et al., 2013; Krawchuk et al., 2013), making it an attractive candidate for the feedback signal. To test this hypothesis, we used *Fgf4^−/−^* embryos, which cannot make PrE (Movie S7), and introduced increasing amounts of wt ESCs, which provide a localized source of FGF4 and are a proxy for wt epiblast cells (Fig. 7A). Thus, increasing amounts of ESCs would allow us to titer the process of PrE specification.

**Figure 7.**
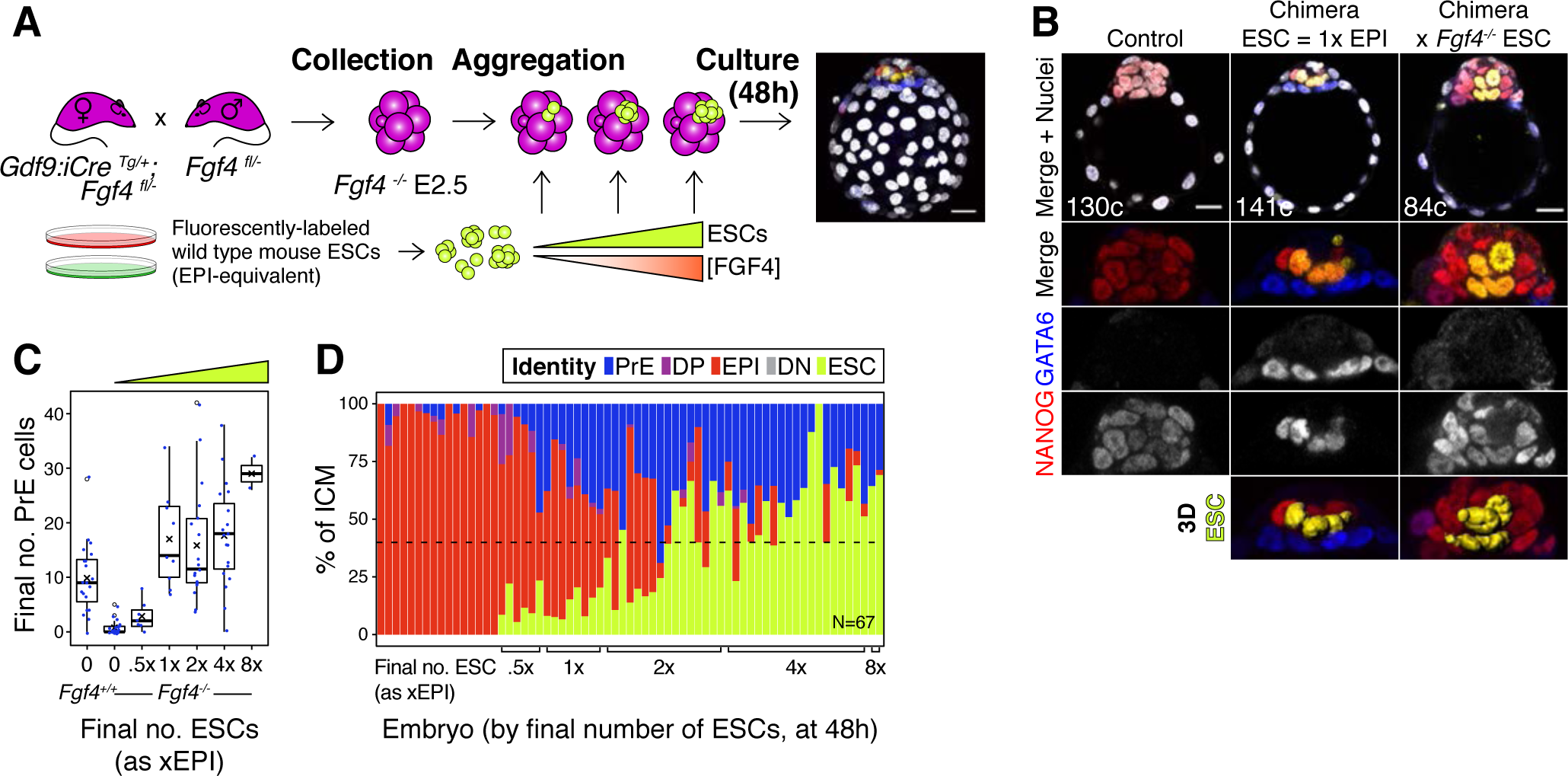
FGF4 provides the dynamic readout of lineage size that determines cell fate specification. **(A)** Experimental design. Maternal zygotic (mz) *Fgf4^−/−^* Embryos were recovered from *Gdf9^iCre/+^; Fgf4^fl/-^* females crossed with *Fgf4^fl/-^* males at the 8-cell stage (2.5 days post-fertilization). Embryos were aggregated with clumps of fluorescently labelled wild type ESCs and cultured for 48-56h, until the late blastocyst stage (equivalent to ∼4.5 days post-fertilization). Control embryos were allowed to develop without adding ESCs. Both chimeric and non-chimeric control embryos were fixed at the end of the culture period and labelled with lineage markers to assess ICM composition. **(B)** Optical cross sections through representative chimeras carrying either wild type or *Fgf4^−/−^* fluorescently labelled ESCs (as indicated) and non-chimeric control embryos labelled for NANOG and GATA6 to identify all ICM cell types. The progeny of the introduced ESCs is labelled in yellow. Total cell count is indicated for each embryo. Lower panels show magnifications of the ICM, with all markers overlaid and for each individual marker in grayscale. Surface renders of ESC compartment within the ICM are shown below. **(C)** Box plots showing absolute number of PrE cells after 48h in wild type, control embryos (no ESCs), *Fgf4^−/−^* control embryos (no ESCs) and *Fgf4^−/−^* chimeric embryos, grouped by the size of the ESC compartment, as in Figure 3. **(D)** Stacked bar plot showing the relative ICM composition in individual embryos (controls or chimeras). Each bar represents the ICM of one embryo and bars are arranged by absolute number of ESCs present in the embryo. Brackets on x-axis indicate the number of ESCs in those embryos, relative to the size of the average wt control epiblast (xEPI), as in (C). Color coding is indicated. All optical cross sections are 5µm maximum intensity projections. In all box plots whiskers span 1.5x the inter quartile range (IQR) and open circles represent outliers (values beyond 1.5x IQR). Cross indicates the arithmetic mean and each dot represents one embryo. Yellow wedges represent the increasing amount of ESCs in each group. PrE: Primitive Endoderm, DP: Double Positive (for NANOG and GATA6), EPI: Epiblast, DN: Double Negative (for NANOG and GATA6), ESC: embryonic stem cell. Scale bars = 20µm.

We found that wt ESCs can rescue PrE specification in *Fgf4^−/−^* embryos (Fig. 7B; S8A; Movie S8). Conversely, *Fgf4^−/−^* ESCs produced chimeras without PrE, phenocopying *Fgf4* null embryos (Fig. 7B; S8A; Movie S9) and confirming FGF4 as the ligand responsible for the rescue. wt ESCs contributed to chimeras with *Fgf4^−/−^* and wt host embryos comparably (Fig. S8B; Fig. S4A) and, although we observed more variation in the size of the host-derived ICM compartment in wt ESC ↔ *Fgf4^−/−^* embryo chimeras than in wt ESC ↔ wt embryo chimeras, this did not relate to the number of ESCs present (Fig. S8C; Fig. 3D). However, the number of wt ESCs present in chimeras dictated the size of the PrE (Fig. 7C). Chimeras with a number of ESCs equivalent to a wt epiblast (1xEPI) showed a full rescue of absolute PrE size (Fig. 7C). Both the absolute and relative size of the PrE in these chimeras depended on the size of the ESC compartment (Fig. 7C-D), with chimeras frequently exhibiting a complete rescue of ICM composition, equivalent to wt (Fig. 7D, dashed line). Of note, chimeras with a high number of ESCs (4x-8xEPI) showed a deviation from wt ICM composition comparable to that observed in wt ESC ↔ wt embryo chimeras (Fig. 7D and 3E; S8D,E and S4D,E). In both cases, ICM composition was disrupted with very high numbers of ESCs (4-8xEPI, Fig. 7D; 3E), likely due to a lack of compensatory proliferation.

Lastly, we tested the need for progenitor cells to rescue the PrE by introducing wild type ESCs into *Fgf4^−/−^* blastocysts, which have totally or partially lost the DP compartment (Fig. S8G, H) (Kang et al., 2013; Ohnishi et al., 2014). Addition of ESCs at this stage failed to rescue the ICM composition in most cases, with only some chimeras specifying low numbers of PrE cells (Fig. S8H,I; Movie S10-11). Taken together, these data demonstrate that FGF4 provides the feedback that couples lineage size and specification, acting as the cell-counting mechanism and eliciting an effect only on uncommitted progenitor cells. Such feedback control enables the system to dynamically adjust the differentiation rate of progenitors in response to perturbations in lineage specified cells to ensure a robust and consistent developmental outcome.

## Discussion

Preimplantation mammalian embryos face two major challenges: to generate sufficient numbers of cells of each of their constituent lineages and to do so before implantation into the uterus. Blastocyst lineages scale with embryo size (Papaioannou and Ebert, 1995; Saiz et al., 2016b), and perturbations in absolute cell numbers are compatible with development to term (Mintz, 1967; Papaioannou et al., 1989; Tarkowski, 1959; 1961), suggesting the variable under control at preimplantation stages is relative, not absolute, lineage size. In this study we show both theoretically and empirically that growth factor-mediated feedback couples cell fate decisions with lineage size in the mouse blastocyst to ensure consistent cell type proportions. Feedback control is widely used in complex systems to buffer noise and ensure robust behavior – in quorum sensing, it alters gene expression to coordinate cellular behaviors at the population level, from bacteria to mammals (Balázsi et al., 2011; Chen et al., 2015; Lander, 2011). Our data show that, in the mammalian blastocyst, growth factor mediated feedback ensures robust patterning independent of absolute embryo size, and it enables regeneration after injury.

Existing models of cell fate specification in the blastocyst have combined a switch mediated by transcription factors (Huang et al., 2007) with growth factor feedback (Bessonnard et al., 2014; Nissen et al., 2017; Schröter et al., 2015; Tosenberger et al., 2017) (reviewed in Simon et al., 2018; Tosenberger et al., 2019). However, the lack of experimental evidence for direct mutual inhibition between the transcription factors NANOG and GATA6 in the embryo, and the non-cell autonomous nature of this cell fate decision (Fig. 1), suggest that intercellular feedback alone could drive this process. In agreement with this expectation, we show that a minimal model containing only indirect mutual inhibition via growth factor signaling is sufficient to (i) robustly generate two ICM compartments (epiblast and PrE), (ii) enable lineage scaling with embryo size, and (iii) adjust for changes in lineage composition. In our model, each cell fate indirectly represses its own specification by promoting the differentiation of neighboring progenitors toward the opposite fate, thus ensuring a balanced cell type composition. This behavior is consistent with the observation that embryos with defective activation of the FGF4-MAPK cascade have an excess of epiblast cells (Brewer et al., 2015; Chazaud et al., 2006; Kang et al., 2013; 2017; Krawchuk et al., 2013; Molotkov et al., 2017; Nichols et al., 2009) and vice versa (Yamanaka et al., 2010).

Although FGF4 signaling controls lineage size in the ICM of the mouse blastocyst, our model is agnostic to the nature of the growth factor involved and could be readily applied to binary cell fate decisions in other contexts. Notably, cell fate proportions during *Dictyostelium discoideum* development are controlled through the secreted factor DIF-1 in an analogous manner (Kay and Thompson, 2001), and members of the TGF-β family mediate negative feedback during skeletal muscle and olfactory epithelium specification in the mouse (McPherron et al., 1997; Wu et al., 2003). In the blastocyst of other mammalian species, NANOG and GATA6 are involved in this fate decision, but the requirement for FGF signaling is less clear (Kuijk et al., 2012; Piliszek et al., 2017; Roode et al., 2012; Soszyńska et al., 2019), suggesting a role for other signaling pathways, or a cell autonomous fate decision. Comparing the prediction of our model, others with alternative configurations, and the result of experimental perturbations, should help elucidate the relative contribution of intercellular signaling and transcription factor networks to lineage specification in these contexts.

The regulative ability of the mouse blastocyst has been extensively tested (Gardner, 1968; Krupa et al., 2014; Mintz, 1967; Tarkowski, 1959; 1961). Introduction of ESCs into morula-stage embryos is used to generate mice entirely derived from ESCs (Lallemand and Brûlet, 1990; Nagy et al., 1990; Poueymirou et al., 2006; Tokunaga and Tsunoda, 1992), which bias host cells toward extra-embryonic lineages (Humięcka et al., 2016). Contrary to what we observe here, introduction of ESCs also affected the host contribution to the ICM (Humięcka et al., 2016). This discrepancy could be due to differences in the time points analyzed, the absence of physical constrains imposed by the zona pellucida in our experiments and/or the higher number of cells introduced by Humięcka and colleagues. We have proposed that a critical element in the regulative ability of the blastocyst stage embryo is the asynchrony in progenitor specification toward epiblast or PrE (Saiz et al., 2016b). Cell cycle phase has been linked to fate decisions in pluripotent stem cells (Pauklin and Vallier, 2013) and it has been shown that heterogeneity in cell cycle phase ensures a robust cell-type composition in *Dictyostelium* (Gruenheit et al., 2018). In the blastocyst, asynchrony in cell cycle phase might enable a dynamic response to changes in growth factor levels, whereby only the subset of progenitors competent to differentiate at any given point will respond to the perturbation. This asynchrony in the cell cycle may ultimately underlie both the progressive allocation of cell fates observed in the ICM and the ability of the system to respond to perturbations in lineage composition.

We probed the robustness of this system by introducing lineage-restricted cells into the embryo to generate chimeras, or by decreasing lineage size using laser cell ablation. Our mathematical model exhibits attractor dynamics through which progenitor cells differentiating after the perturbation would adopt the fate needed to restore the lineage balance, a prediction corroborated by our experimental perturbations. A key element in this response is the fact that NANOG levels in a cell are inhibited by those in its neighbors through a lateral inhibition mechanism. Such growth factor-mediated lateral inhibition can explain the salt-and-pepper distribution of cell types in the ICM (Chazaud et al., 2006; Rossant et al., 2003), although cell movement within the ICM likely precludes the observation of the canonical, alternate distribution of fates. Besides Delta-Notch signaling, lateral inhibition can also result from mechanical cues (Xia et al., 2019) and from secreted signaling molecules (Thompson et al., 2004). It has been proposed that high local concentrations of FGF4 may underlie this stochastic distribution of fates in the ICM of the blastocyst (Bessonnard et al., 2014; Kang et al., 2013; 2017; Tosenberger et al., 2017) and that the identity of neighbor cells affects fate choice (Fischer et al., 2017). In agreement with this view, our proposed configuration of the regulatory network leads to spontaneous divergence of fates among neighboring cells: high availability of growth factor around cells with high NANOG levels (FGF4 producers (Frankenberg et al., 2011; Guo et al., 2010; Nowotschin et al., 2019; Ohnishi et al., 2014)) induces NANOG downregulation and PrE fate among surrounding progenitors. Internalization of ligand-receptor complexes by receiving cells, differential intracellular feedback and the transient tandem expression of FGF receptors 1 and 2 in cells adopting a PrE fate (Kang et al., 2017; Molotkov et al., 2017; Nowotschin et al., 2019; Ohnishi et al., 2014) may further contribute to the directionality of this local gradient and the resulting induction of opposite fates in neighboring cells.

Further support for the role of FGF4 in the epiblast/PrE decision comes from the finding that introduction of wt ESCs can rescue the all-epiblast ICM found in *Fgf4^−/−^* embryos (Fig. 7D). Treatment of *Fgf4^−/−^* embryos with saturating doses of recombinant FGF4 results in differentiation of all ICM cells toward PrE, instead of a normal balance of epiblast and PrE cells (Kang et al., 2013; Krawchuk et al., 2013), presumably due to homogeneous exposure to high levels of ligand by all progenitor cells. In our chimeras, however, the ESCs introduced effectively act as a wt epiblast and provide a local source of FGF4. Consequently, the size of the PrE population is dictated by the amount of ESCs present (Fig. 7C), suggesting that only progenitors neighboring ESCs are exposed to the signal and adopt a PrE fate. Although our current data lacks the spatial resolution to determine the relative position of emerging PrE cells, these experiments establish a direct link between both fates via FGF4 and serve as a proxy for PrE specification in a wt context.

Our study uncovers how cell fate choice and lineage size are coupled via growth factor signaling to ensure robust patterning and morphogenesis in a self-organizing developmental system – independently of size, and without the need for morphogen gradients. These findings provide a framework for our current understanding of signaling and cell fate decisions in the early mammalian embryo and may be generalizable to the formation of other autonomous developmental units.

## Materials and Methods

### Mouse strains and husbandry

All animal work was approved by Memorial Sloan Kettering Cancer Center’s Institutional Animal Care and Use Committee. Animals were housed in a specific pathogen-free facility under a 12h light cycle (6am-6pm). Embryos were obtained from natural matings of 4-12 week-old females to stud males. Alleles used and their original source are summarized in Table 1. All mice were maintained on a mixed genetic background.

### Embryo staging

Embryonic days (E) were determined by considering noon of the day of detection of the copulation vaginal plug as E0.5. Higher-resolution staging based on total cell number was used throughout as previously (Kang et al., 2013; Plusa et al., 2008). To determine the initial developmental stage in experiments involving *in vitro* culture of blastocysts without a ubiquitous nuclear reporter, a subset of embryos from each litter (2-4 embryos) were fixed at the time of collection, as a reference for the developmental stage of the entire litter (referred to as Littermates in data tables) as previously (Grabarek et al., 2012; Saiz et al., 2016b). In all experiments, individual litters were treated as experimental units to reduce variability due to stage differences between litters and to control for batch effects.

### Embryo recovery and handling

Embryos were flushed from dissected oviducts (8-16 cell stage) or uterine horns (blastocyst stages) using forcing Flushing and Holding Media (FHM, Millipore), as previously described (Behringer et al., 2014). Live embryos were manipulated in FHM. All solutions used to handle or recover live embryos were pre-warmed at 37°C. Whenever not being handled under the microscope, embryos were kept at 37°C in a dry incubator box. Zona pellucidae were removed from embryos by brief washes in Acid Tyrode’s solution (Millipore) (Behringer et al., 2014) and returned to FHM as soon as the zona dissolved. Blastocysts were fixed in a solution of 4% paraformaldehyde (PFA, Bio-Rad) in phosphate buffered saline (PBS) for 10 minutes at room temperature and preserved in PBS at 4°C.

### Embryo genotyping

Embryos were lysed for genotyping in 10µl of lysis buffer (10mM Tris, pH 8.5, 50mM KCl, 0.01% gelatin and 300µg/ml of Proteinase K) for 50 minutes at 50°C, followed by 10 minutes at 90°C to inactivate Proteinase K (Artus et al., 2005). 2µl of the lysate was used for genotyping (primer sequences indicated in Table 2).

### Embryo culture and live imaging

Embryos were cultured on 35mm Petri dishes (Falcon) within microdrops of 10-15µl of Potassium Simplex Optimized Media with amino acids (KSOM-AA, Millipore) under mineral oil (Sigma), at 37°C, in a humidified 5% CO2 atmosphere. Prior to culture, embryos were rinsed 3x in drops of KSOM-AA. For live imaging, embryos were cultured on 35mm glass-bottom dishes (MatTek) within 5-10µl drops of KSOM-AA.

Images of live embryos were acquired using Zeiss LSM880 laser-scanning confocal microscopes. In cell ablation experiments, images were acquired using a Zeiss C-Apochromat 40x/NA1.1/WD0.21mm objective. For all other experiments, images were acquired using a Zeiss EC Plan-Neofluar 40x/NA1.3/WD0.17mm. GFP was excited using a 488nm Argon laser at 20µW. mKate2 was excited using a 543nm HeNe laser or a 561nm DPSS 561-10 laser at 90µW. Laser power was measured through a Zeiss Plan-Neofluar 10x/NA0.3 objective prior to each imaging session with a light meter (Coherent) and the laser output adjusted to match laser power across experiments. 80µm stacks were acquired through embryos, at 2µm intervals, every 15 min. Although these stacks do not capture an entire blastocyst, they encompass the ICM while limiting laser exposure to about 30-40s per time point and embryo. Time lapse movies were 16-20h long.

### Cell culture

Embryonic Stem Cells (ESCs) used were *CAG:H2B-EGFP^Tg/+^* (Hadjantonakis and Papaioannou, 2004) and *Hex^Venus/+^*; *CAG:H2B-tdTomato^Tg/+^* (Morgani et al., 2013). ESCs were cultured feeder-free on 0.1% gelatin-coated tissue culture dishes (Falcon) in DMEM supplemented with 2mM L-glutamine, 0.1mM MEM non-essential amino acids (NEAA), 1mM sodium pyruvate, 100 U/ml penicillin, 100µg/ml streptomycin (all from Life Technologies), 0.1mM 2-mercaptoethanol (Sigma), 10% Fetal Bovine Serum (Hyclone) and 1000 U/ml of recombinant leukemia inhibitory factor (LIF). Cells were passaged by brief incubation at 37° C in Trypsin-EDTA (0.25% or 0.05%, Life Technologies), neutralization with Serum/LIF media, centrifugation and resuspension in the desired culture media before replating.

### Embryo-embryo aggregation chimeras

Embryos were isolated in the morning of the third day of development (E2.5) and those with 4 or fewer cells were discarded. Morulae (8-16 cells) from CD1 x *CAG:H2B-EGFP^Tg/+^* (Hadjantonakis and Papaioannou, 2004) crosses were sorted into GFP+ and GFP−. GFP+ embryos (*CAG:H2B-EGFP^Tg/+^*) were used in the generation of both wt ↔ GFP+ and *Gata6^−/−^* ↔ GFP+ chimeras (Fig. 1B), whereas GFP-wild type (wt) morulae were either used for wt ↔ GFP+ chimeras or cultured as un-manipulated controls. For *Gata6^−/−^* ↔ GFP+ chimeras, the *Gata6^−/−^* component was generated as in Figure 1D (see Table 1 for alleles). The use of *Gdf9^iCre^* (Lan et al., 2004) and a floxed *Gata6* (Sodhi et al., 2006) gave greater numbers (>25%) of *Gata6^−/−^* embryos than would result from standard heterozygous *inter se* crosses.

After removal of the zona pellucida, uncompacted, 8-cell stage embryos, were disaggregated and blastomeres reaggregated in the desired proportions, as described elsewhere (Behringer et al., 2014; Eakin and Hadjantonakis, 2006; Mintz, 1964; Tarkowski and Wróblewska, 1967) and summarized in Fig. 1A, D. mz*Gata6^−/−^* embryos are not always null and can show mosaicism. Therefore, in all *Gata6^−/−^* ↔ GFP+ chimeras the *Gata6^−/−^* complement was clonal. All *Gata6^−/−^* ↔ GFP+ chimeras were genotyped retrospectively for the presence of both the conditional and null *Gata6* alleles, and phenotyped based on the presence/absence of GATA6 in immunofluorescence images.

### ESC-embryo chimeras

ESCs and aggregation dishes were prepared as described in (Behringer et al., 2014) and denuded E2.5 embryos (8-16 cells) of the desired genotype were placed within the indentations on the culture dish. ESCs were rinsed twice in KSOM-AA to dilute the FBS and LIF present in the cell media and clumps of the desired number of ESCs placed in contact with each embryo (see Fig. S4A, S8B for cell numbers). Aggregates were then allowed to develop *in vitro* for 48-56h (Fig. 3A), until the late blastocyst stage, when they were fixed in 4% PFA.

For blastocyst ESC injections, a PrimeTech Piezo drive (Sutter Instruments) attached to an Eppendorf CellTram microinjectiong system was used to assist with injections. Individual embryos were held with a holding pipette (VacuTip, Eppendorf) by the ICM end, while a Piezo Drill flat tip needle (Eppendorf) was used to introduce ESCs into the blastocyst cavity (Fig. S8G). After injection, each litter was allowed to develop for 28-31h in KSOM-AA. 3-4h after injection, embryos were denuded to allow for cavity expansion while maintaining blastocyst morphology. Reference littermates were fixed in 4% PFA right after injection into the rest of the embryos.

### Laser cell ablation

Ablation experiments are summarized in Figure 5A and C. To identify PrE cells, we used a nuclear reporter for *Pdgfra* expression (*Pdgfra^H2B-GFP/+^* (Hamilton et al., 2003; Plusa et al., 2008)) (Fig. S5A). To visualize epiblast cells, we combined it with a spectrally-distinct, ubiquitous nuclear mKate2 reporter (*ROSA26:CAG:3x-nls-mKate2^Tg/Tg^*, henceforth referred to as *R26:mKate^Tg/Tg^*) (Susaki et al., 2014), in which epiblast cells are labelled by nuclear mKate2, but not GFP (Fig. S5A,B). For ablation of PrE cells, most embryos used were obtained from CD1 females intercrossed with *Pdgfra^H2B-GFP/+^* stud males (Fig. 5A). For ablation of epiblast cells (and some PrE ablation experiments) all embryos used were from intercrosses of *R26:mKate^Tg/Tg^* females and *Pdgfra^H2B-GFP/+^*; *R26:mKate^Tg/Tg^* males (Fig. 5C).

Embryos were collected at sequential time points between noon and late evening of the fourth day of development (∼E3.5 to E4.0) to cover the period of PrE and epiblast specification (∼30 to 110 cells, Fig. S5E). To determine the number of PrE or epiblast cells in each embryo (and the total cell number in embryos expressing mKate2), a z-stack through each embryo was acquired prior to cell ablation (Fig. 5A, C). Nuclei were considered to belong to the PrE if the GFP signal from the *Pdgfra^H2B-GFP^* allele was markedly higher than in their neighbors on each optical plane (Fig. 5b; S5a). Nuclei with intermediate levels of GFP were scored as putative progenitor cells, whereas nuclei with no GFP (but labelled with mKate2) were scored as epiblast (Fig. S5A, B, see Fig. S5D-E for estimates of each population size in live embryos compared to fixed samples stained for molecular markers). Using time-lapse imaging, we confirmed that cells classified as PrE maintained or upregulated *Pdgfra* expression as they developed, whereas epiblast cells maintained low levels (Fig. S5F, see Data Processing, below, for details on cell classification). On the other hand, cells classified as progenitors upregulated or downregulated *Pdgfra* expression over time, as they adopted PrE or epiblast identity, respectively (Fig. S5F).

Cells were eliminated by repeatedly focusing the beams of an 800-nm Ti:Sapphire femtosecond laser at 10-12% output (Coherent), onto the central region of each nucleus to be ablated (approximately 30-50% of the nuclear area in the section). The number of pulses was determined empirically on a test embryo for each experiment, so that a clear wound was observed on the fluorescent nucleus, but no obvious damage was inflicted to the neighboring cells (typically between 125-250x iterations) (Fig. 5B, D). Although target nuclei were selected at random throughout the ICM, when possible, nuclei located on the same z plane were targeted, to minimize the length of the procedure. Intact control embryos were only subject to the initial imaging step to estimate the size of each ICM population. Random control embryos were *R26:mKate^Tg/Tg^* embryos (not carrying the *Pdgfra^H2B-GFP^* allele) in which a number of ICM cells equivalent to the total number of PrE or epiblast cells found in GFP+ littermates (100% PrE- or epiblast-equivalent) was randomly selected and ablated using the same settings.

After cell ablation, live intact and targeted embryos, were imaged for 16-20h, as described above. Laser ablation generated a visible wound in nuclei expressing H2B-GFP, which allowed tracking of targeted and intact cells and assessing nuclear fragmentation as a mark of cell death (Fig. 5E; Movie S4). However, mKate2 was extinguished after repeated laser illumination and consequently targeted epiblast cells generally could not be tracked. After time-lapse imaging, embryos were allowed to develop further in an incubator, until a total of 24h after the time of collection, before being fixed individually in 4% PFA.

### Immunofluorescence

Whole-mount embryo immunofluorescence was performed as described previously (Saiz et al., 2016a; 2016b). Primary antibodies and the dilutions used are provided in Table 3. Secondary antibodies were from Life Technologies, except the AlexaFluor®488 anti-chicken, which was from Jackson ImmunoResearch.

### Image acquisition of fixed samples

Immunolabeled embryos were mounted on 35mm glass-bottom dishes (MatTek), within micro drops of a 5µg/ml solution of Hoechst 33342 in PBS and imaged using a Zeiss LSM880 laser-scanning confocal microscope, equipped with an oil-immersion Zeiss EC Plan-Neofluar 40x/NA1.3/WD0.17mm. Z-stacks were acquired through whole embryos with 1µm step between optical slices. Laser power was measured for each laser line prior to each imaging session and parameters adjusted so as to keep laser power consistent for each primary-secondary antibody combination across experiments over time – except for the 405 channel, which was used to excite the nuclear label (Hoechst 33342), and was solely used for image segmentation.

### Image processing

Nuclear image segmentation of still images and manual image correction was performed using the MINS software (Lou et al., 2014) as previously described (Morgani et al., 2018b; Saiz et al., 2016a). MINS is freely available at https://github.com/therealkatlab/MINS (requires MATLAB license). Missing nuclei, or multiple nuclei segmented as one (under-segmentation) were measured manually using ImageJ (Rasband, W.S., ImageJ, U. S. National Institutes of Health, Bethesda, Maryland, USA, https://imagej.nih.gov/ij/, 1997-2018) and the measured values for each channel, as well as XYZ coordinates, added to the data table.

Time lapse images were processed using Imaris (Oxford Instruments). Individual ICM cells were identified based on the presence of H2B-GFP and/or nuclear mKate2 and tracked manually over the course of the movie using the spots function. Cell death or mitotic events were labelled as such for each individual cell (Fig. S5G). GFP levels are a proxy for *Pdgfra* expression and allow the classification of cell types over time (Fig. S5F, G). Cell identity was assigned visually based on GFP levels at the time of the experiment (Fig. S5A, E) and verified retrospectively (and reclassified when necessary) after quantification of GFP levels.

### Data processing

Fluorescence data obtained after segmentation with MINS and manual curation was processed as described in (Morgani et al., 2018b; Saiz et al., 2016b), with certain modifications for subsets of the data. For samples where GFP fluorescence was detected indirectly with an anti-GFP antibody (Fig. 1; S2), anti-GFP::AF®488 fluorescence was log-transformed and corrected for fluorescence decay along the Z axis by fitting a linear model, as described in (Saiz et al., 2016a). For all other samples (live or fixed), an Empirical Bayesian slope correction step was applied to log-transformed, Z-corrected data, as in (Saiz et al., 2016b). Detailed descriptions of the correction steps can be found in https://github.com/nestorsaiz/saiz-et-al_2020/tree/master/notebooks

Two different antibodies were used to detect NANOG and GATA6 expression (Table 3). Rabbit anti-NANOG (NANOG(rb)) and goat anti-GATA6 (GATA6(gt)) were used for most samples, as previously (Kang et al., 2017; Morgani et al., 2018b; Saiz et al., 2016b; Schrode et al., 2014). When a rat anti-NANOG (NANOG(rat)) or a rabbit anti-GATA6 (GATA6(rb)) were used (Table 3), fluorescence values were transformed to NANOG(rb)- and GATA6(gt)-equivalents, respectively, using a linear regression model generated from samples stained with two antibodies for each marker. A more detailed description can be found in https://github.com/nestorsaiz/saiz-et-al_2020/blob/master/notebooks/nanogata-tx.ipynb

Cell identity was assigned to ICM cells based on relative NANOG and GATA6 expression, re-scaled against their maxima in each litter to account for variability between litters and over time. When separated based on NANOG and GATA6 levels, ICM cells typically form four clusters, corresponding to the epiblast (NANOG+), PrE (GATA6+), double positive (DP, progenitor cells) and NANOG-low or NANOG-epiblast (EPI-lo: GATA6-, NANOG-) (Kang et al., 2017; Morgani et al., 2018b; see Saiz et al., 2016b). To automatically identify cell populations in the ICM we implemented hierarchical clustering. We determined empirically that the agglomerative UPGMA (unweighted pair method with arithmetic mean) captured the PrE and epiblast clusters comparably to k-means clustering (as used in (Kang et al., 2017; Morgani et al., 2018b; Saiz et al., 2016b)) across all blastocyst stages, however, it outperformed k-means when classifying DP and NANOG-low epiblast cells. More detailed descriptions of the transformation and classification steps can be found in https://github.com/nestorsaiz/saiz-et-al_2020/tree/master/notebooks

Corrected fluorescence values of H2B-GFP (*Pdgfra* expression) for ICM cells were used to determine changes in cell identity over time in time-lapse movies. To reduce noise in the data, a moving average was calculated for each cell, with a window of 4 timeframes (1h). A simple classifier was thus devised to assign cell identity automatically to individual cells based on these GFP levels. Thresholds for fluorescence were manually determined for each litter analyzed, both at the start and the end of the movie, which were used to determine a threshold slope for PrE and epiblast identities. The classifier followed two rules: (1) cells classified as PrE or epiblast maintain that identity for the remainder of the movie – based on (Saiz et al., 2016b; Xenopoulos et al., 2015) and the lack of oscillation on GFP levels or obvious, systematic shifts from high to low levels, or vice-versa (Fig. S5F, G) – (2) cells classified as DP at t=0 become PrE or epiblast once their level of GFP remains above or below the respective thresholds for at least 2h – to account for fluctuations and noise in data (Fig. S5G).

### Data and code availability

All data processing was done in R version 3.4.2, using RStudio as an interactive development environment. All processed data as well as the code used to transform data and classify cells is publicly available at https://github.com/nestorsaiz/saiz-et-al_2020 and upon request. All raw confocal images and data tables will be freely available on Figshare with DOI <u>10.6084/m9.figshare.c.4736507</u>.

## Acknowledgements

We thank Drs Frederic Geissmann, Pierre-Louis Loyher, Alison North and Christina Pyrgaki for access to their two-photon systems and for invaluable help in setting up the experimental conditions for cell ablation in embryos; Dr Pavak Shah for advice on laser ablation in different contexts; Drs Jennifer Nichols and Stanley Strawbridge for discussions of complementary projects; Drs Alfonso Martinez Arias, Berenika Plusa, Pedro Rocha, Christian Schröter, Eric Siggia, Philippe Soriano, Joana Vidigal, and members of the Hadjantonakis lab for insightful discussions and critical feedback. The authors would like to dedicate this work to the memory of Prof Andrzej K Tarkowski and Dr Yoshiki Sasai, whose pioneering work has long been a source of inspiration.

## Author contributions

NS and AKH conceived and planned the experiments. NS performed all embryo experiments. LMB and JGO built the mathematical model and performed *in silico* experiments. NS, SR, HG and JPH manually curated immunofluorescence data. NS and SR processed time-lapse data. SR and JPH assisted with sample genotyping. NS designed and wrote data processing pipelines and analyzed experimental data with assistance from SR and HG. NS and JGO generated graphs and figures. NS, JGO and AKH wrote the manuscript with input and approval from all authors.

## SUPPLEMENTARY TEXT

### 1 Molecular model

The minimal model used in this paper can be derived from the following kinetic model describing the interactions between Nanog (*N*), Gata6 (*G*) and FGF (*F*) shown in Fig. S1:

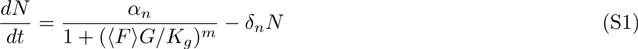

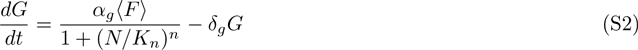

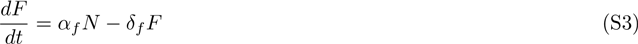

Here Nanog and Gata6 in a given cell inhibit each other indirectly, through FGF signaling coming from the cell itself and from its neighbors (represented by the average term ⟨*F*⟩ in Eqs. S1 and S2). The average FGF term, ⟨*F*⟩, enters the two inhibition terms asymmetrically: in the inhibition of Nanog by Gata6 (first term in the right-hand side of Eq. S1), FGF affects directly the repressor concentration and appears in the denominator, whereas in the inhibition of Gata6 by Nanog (Eq. S2), FGF modulates the repression term globally and appears in the numerator. This corresponds to a situation in which FGF activates ERK (which in turn activates Gata6) *after the repression by Nanog has already taken place* (via negative modulation of the receptors FGFR1 and FGFR2), leading to a global modulation of the repressed Hill function, as shown in Eq. (S2). On the other hand, ERK is considered to inhibit Nanog, so that the effect of FGF on Nanog can be modeled as *affecting directly its repressor* (here considered to be Gata6 itself), thereby multiplying the concentration of that repressor in the denominator of the inhibitory Hill function of Eq. (S1). We note, however, that the minimal model obtained below could also represent other molecular architectures of the Nanog-Gata6 circuit, and thus can be interpreted as a general representation of a growth-factor-mediated mutual inhibition.

**Figure S1:**
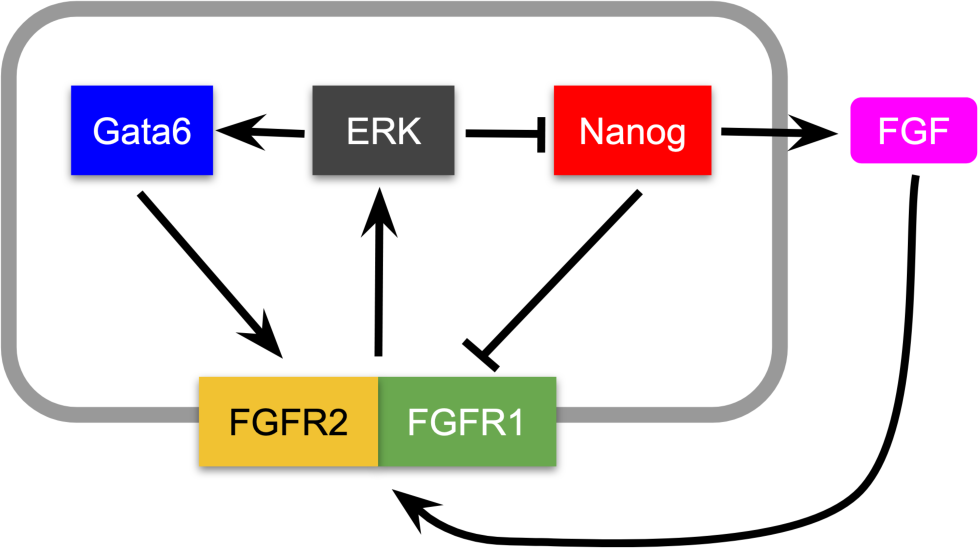
Potential molecular mechanism underlying the Epi-PrE decision.

### 2 Reduction to one variable

We now assume that Gata6 and FGF are faster than Nanog, so that Eqs. (S2) and (S3) can be considered to be in quasi-steady state. We thus can make the right-hand side of those equations equal to zero, and substitute the resulting expressions of *G* and *F* into Eq. (S3). Nondimensionalization and a minor algebraic rearrangement of the production term lead to a single ODE per cell:

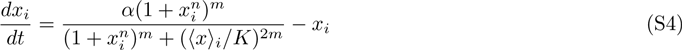

where *x_i_* represents the concentration of Nanog in cell *i* measured in units of *K_n_* and time is measured in units of 1*/*δ*_n_*. The signaling term ⟨*x*⟩*_i_* denotes the average value of *x* in the neighborhood of cell *i*, including *x_i_* itself, with weights that can be fixed at will. Here we assume equal weights in the immediate neighborhood –nearest neighbors and the self– and 0 weights for the rest of the cells. The dimensionless parameters *α* and *K* are related with the original parameters of model (S1)-(S3) through the following relationships:

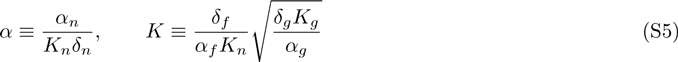

The dimensionless parameters of the model that we use in this paper, together with those of the agent-based model described in what follows, are given in Table S1 below.

**Table S1:**
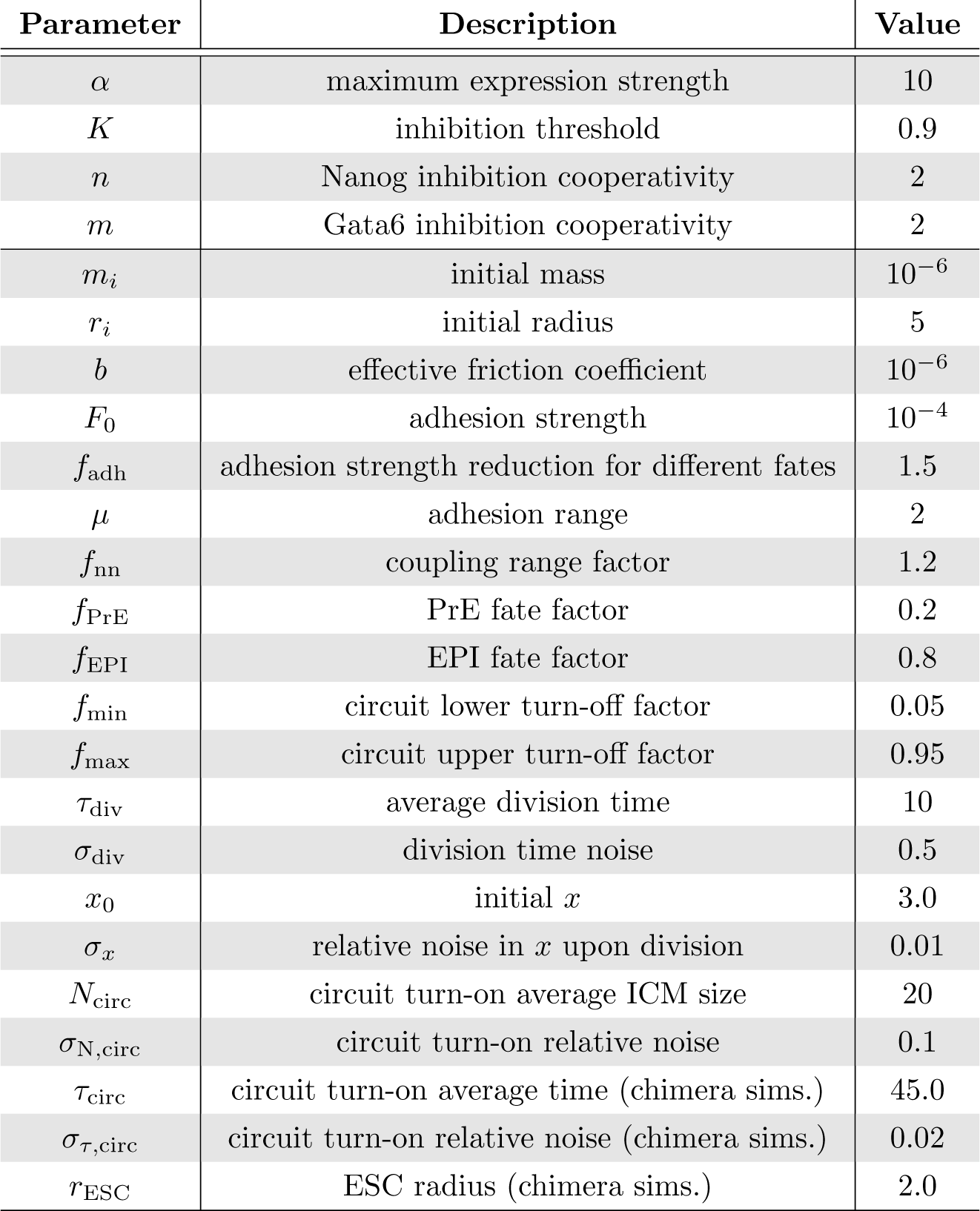
Parameter values. The parameters of the biochemical circuit (top 4 parameters, above the line) are given in dimensionless units. The parameters of the agent-based model (below the line) are given in arbitrary units, with time being rescaled *a posteriori* to match the experimental observations approximately.

| Parameter | Description | Value |
| --- | --- | --- |
| $\alpha$ | maximum expression strength | 10 |
| $K$ | inhibition threshold | 0.9 |
| $n$ | Nanog inhibition cooperativity | 2 |
| $m$ | Gata6 inhibition cooperativity | 2 |
| $m_i$ | initial mass | $10^{-6}$ |
| $r_i$ | initial radius | 5 |
| $b$ | effective friction coefficient | $10^{-6}$ |
| $F_0$ | adhesion strength | $10^{-4}$ |
| $f_{adh}$ | adhesion strength reduction for different fates | 1.5 |
| $\mu$ | adhesion range | 2 |
| $f_{nn}$ | coupling range factor | 1.2 |
| $f_{PrE}$ | PrE fate factor | 0.2 |
| $f_{EPI}$ | EPI fate factor | 0.8 |
| $f_{min}$ | circuit lower turn-off factor | 0.05 |
| $f_{max}$ | circuit upper turn-off factor | 0.95 |
| $\tau_{div}$ | average division time | 10 |
| $\sigma_{div}$ | division time noise | 0.5 |
| $x_0$ | initial $x$ | 3.0 |
| $\sigma_x$ | relative noise in $x$ upon division | 0.01 |
| $N_{circ}$ | circuit turn-on average ICM size | 20 |
| $\sigma_{N,circ}$ | circuit turn-on relative noise | 0.1 |
| $\tau_{circ}$ | circuit turn-on average time (chimera sims.) | 45.0 |
| $\sigma_{\tau,circ}$ | circuit turn-on relative noise (chimera sims.) | 0.02 |
| $r_{ESC}$ | ESC radius (chimera sims.) | 2.0 |

### 3 Agent-based modeling

We apply the biochemical model described above to a population of proliferating spherical cells that simulate the developing embryo [1], whose mechanics are simply governed by:

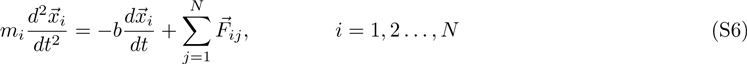

Here 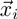 denotes the position of the center of cell *i* with mass *m_i_*, and 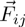 represents the force between cells *i* and *j*, which is repulsive for separations between cell centers smaller than the sum of their radii, *r_ij_* ≡ *r_i_* + *r_j_*, attracting for larger separations up to *µr_ij_*, and zero for separations larger than *µr_ij_*. This is represented by:

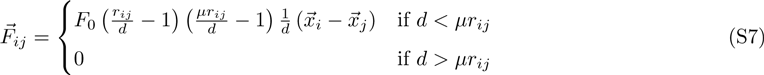

where *d* is the Euclidean distance between the cell centers. The interaction strength *F*_0_ between two cells is made to depend on their relative fates, being smaller between cells of different type than between cells of the same type (see Table S1). In the biochemical circuit described above, Eq. (S4), two cells are considered neighbors if the separation between their centers is smaller than the sum of their radii multiplied by a factor *f*_nn_.

Cellular proliferation is modeled by making each cell divide a certain time after its previous division. We consider that this time varies uniformly in the interval [τ_div_ - σ_div_, τ_div_ + σ_div_]. The daughter cells are placed at symmetrically selected positions around the center of the mother cell along a randomly chosen direction, separated a distance equal to the radius of the mother cell. Upon each division, cells (with the exception of externally added embryonic stem cells, see below) reduce their mass a factor 2 and their radius a factor 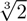, which corresponds to their volume being halved. The daugther cells inherit all other properties from their mother, including their velocity, mass, fate, and Nanog level *x*. Nanog is allowed to vary stochastically upon division, by a uniformly distributed random amount with relative amplitude σ*_x_*.

The biochemical circuit is turned on after the ICM has reached a certain average ICM size *N*_circ_ (with relative noise σ_N,circ_), to account for the fact that experimental observations do not reveal any meaningful dynamics of Nanog nor Gata6 in the first stages of embryonic development. The scaling experiments are modeled by multiplying *N*_circ_ by the corresponding factor (2x, 0.5x, 0.25x). In the chimera simulations, circuit turn-on is applied instead at a specific average time *τ*_circ_ (with relative noise (σ_τ,circ_) that is consistent with the turn-on ICM size exhibited by the wild-type embryos.

The fate of a cell is determined on the basis of the value of the *x* variable in that cell at any given time. Specifically, if *x* is larger than a fraction *f*_EPI_ of its maximum possible value (*x*_M_ = ⍺*/*(1 + 1*/*(2*K*)^2^*^m^*)), the cell is considered to be an EPI cell. In turn, if *x < f*_PrE_*x*_M_ the cell is considered a PrE cell. These conditions are chosen to mimic the way in which cell fates are usually assigned experimentally [2]. Once those fates are reached, the biochemical circuit is turned off. Specifically, *x*(*t*) is only updated following the dynamics described by Eq. (S4) when *f*_min_*x*_M_ *< x < f*_max_*x*_M_. This assumption reflects the fact that our biochemical models ignores downstream processes that follow EPI and PrE specification, making these cell-fate choices irreversible [3]. The limit factors *f*_PrE_ and *f*_min_ (resp. *f*_EPI_ and *f*_max_) are assumed to differ somewhat (see Table S1), in order to make the irreversible character of the decision robust to noise in *x*.

The ablation experiments are modeled by eliminating random sets of cells (within a specific fate if applicable) at specified ICM sizes, and allowing the remaining cells to redistribute according to the dynamics of Eq. (S6). The chimera experiments are modeled by adding cells with a distinct (ESC) fate, which do not obey the biochemical dynamics described by Eq. (S4) and have a fixed radius *r*_ESC_ that does not decrease upon proliferation. The ESCs are added recursively at a location adjacent to the ICM cell whose position is farthest from the center-of-mass of the simulated embryo at that particular time instant, outside of the embryo and separated from it a distance equal to the radius of that ICM cell. As in the case of the ablation, the added ESCs reorganize quickly after addition, following the dynamics of Eq. (S6).

The model parameters used throughout the paper are given in Table S1. Note that the cell-fate decision circuit depends only on two dimensionless parameters, *α* and *K*, plus two Hill coefficients *n* and *m*. The agent-based model provides the substrate on which the circuit operates, effectively mixing the cells as they proliferate, and providing them with a neighborhood that represents the entire population. The choice of parameters for the agent-based simulations thus does not affect the qualitative behavior of the biochemical circuit introduced here.

---

**Supplementary Figure 1 (Related to Figure 1).**
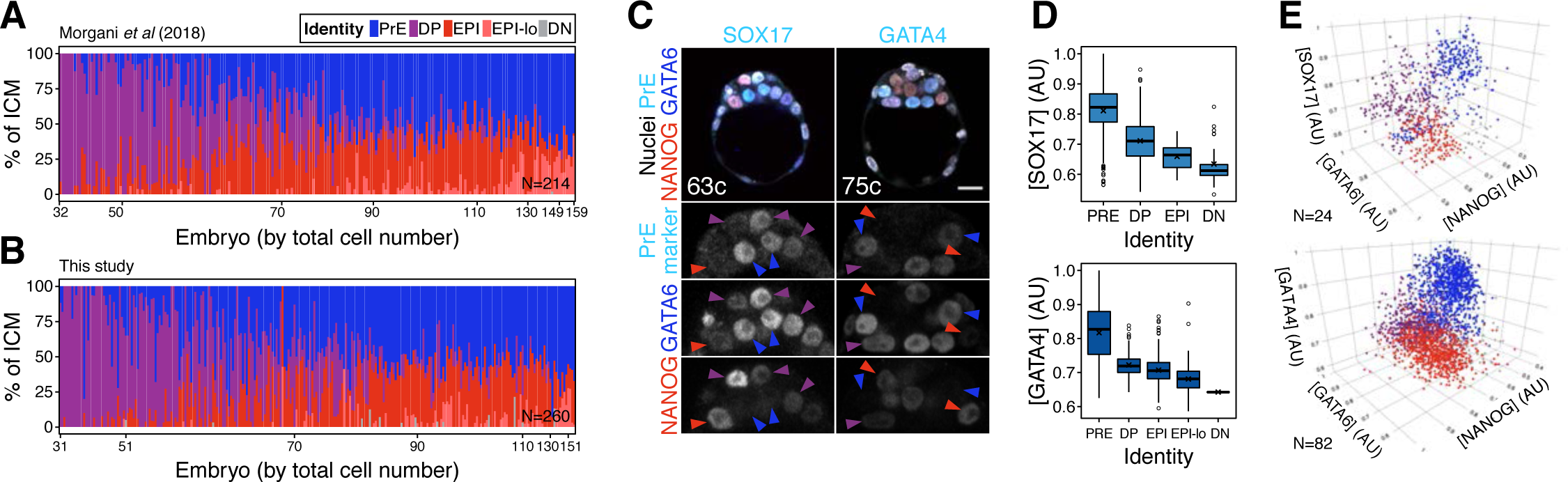
**(A)** Stacked bar plots showing progression of lineage composition in the ICM of wild type blastocysts fixed upon collection analyzed in (Morgani et al., 2018b). ICM cell types were re-assigned using hierarchical clustering, as described in the Methods section. **(B)** Stacked bar plots showing progression of lineage composition in the ICM of wild type blastocysts fixed upon collection generated in this study. Color coding for cell types is indicated. Each bar represents the ICM of a single embryo. Embryos are arranged by ascending total cell number (from 32 up to >150 cells, comprising all the stages of blastocyst development). N indicates number of embryos in the plot. **(C)** Optical cross sections through representative immunofluorescence images of blastocysts stained for NANOG (red), GATA6 (blue) and either SOX17 or GATA4 (cyan), as indicated. Arrowheads point at cells classified as PrE (blue), epiblast (red) or DP (purple). **(D)** Average SOX17 and GATA4 levels in each ICM cell type in a cohort of 24 embryos (SOX17) and 82 embryos (GATA4). Values are natural log re-scaled to 0-1 for each dataset. **(E)** 3D scatterplots for the three markers shown in (C, d). Each spot represents one cell. Spots are color-coded for cell identities as indicated in (A). Cells were classified using only relative NANOG and GATA6 levels, as described in the Methods. In all box plots whiskers span 1.5x the inter quartile range (IQR) and open circles represent outliers (values beyond 1.5x IQR) and cross represents the arithmetic mean. All optical cross sections are 5µm maximum intensity projections. Total cell counts are indicated for each embryo within the merged images. PrE: Primitive Endoderm, DP: Double Positive (for NANOG and GATA6), EPI: Epiblast, EPI-lo: low NANOG epiblast, DN: Double Negative (for NANOG and GATA6). Scale bars = 20µm.

**Supplementary Figure 2 (Related to Figure 1).**
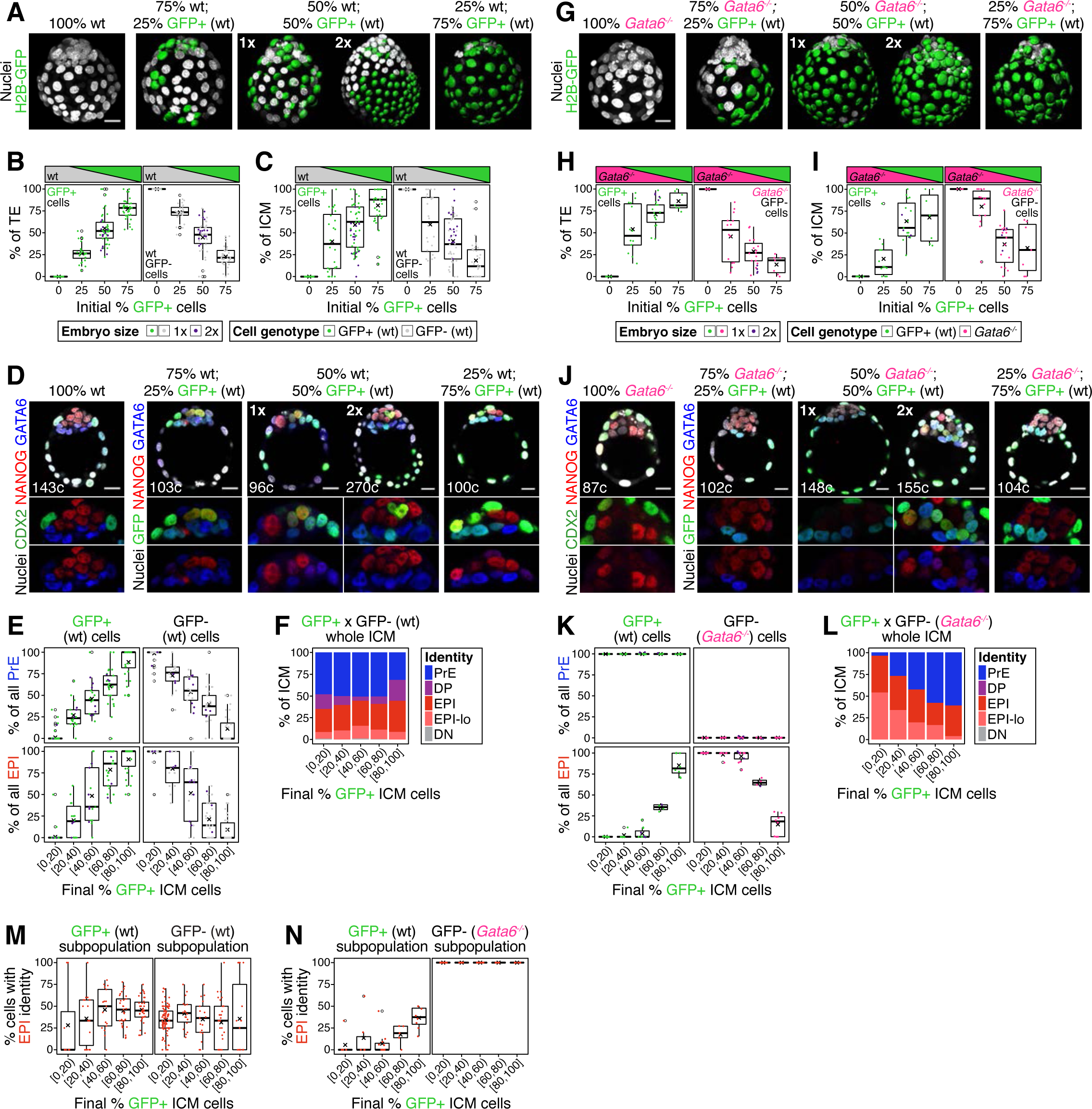
**(A)** Maximum intensity projections of all nuclei of a series of representative wild type control and wt ↔ GFP+ chimeras made with increasing % of wild type GFP+ cells, as described in Fig. 1A. 1x indicates normal size embryo, resulting from aggregating 8 cells, 2x indicates double-sized embryo, resulting from aggregating 2x intact 8-cell stage embryos. **(B)** Box plots showing the relative contribution of GFP+ (green) and GFP−, wild type (gray) cells to the embryo for each of the initial % of GFP+ cells, as shown in (A) Dark green spots represent double embryos (2x). **(C)** Box plots showing the relative contribution of GFP+ (green) and GFP-, wild type cells (gray) to the ICM for each of the initial % of GFP+ cells, as shown in (A). **(D)** Optical cross sections through embryos shown in (A) and immunostained for the markers indicated. **(E)** Box plots showing the relative contribution of GFP+ (green) and GFP-, wild type cells (gray) to each the PrE and the epiblast. Embryos are grouped by the final fraction of GFP+ ICM cells, as indicated on the x-axis **(F)** Stacked bar plots showing the ICM composition in wt ↔ GFP+ chimeras, grouped by the final fraction of GFP+ ICM cells, as in (E) and as indicated on the x-axis. Color coding is indicated. **(G)** Maximum intensity projections of all nuclei of a series of representative *Gata6^−/−^* control and *Gata6^−/−^* ↔ GFP+ chimeras made with increasing % of GFP+ (wt) cells, as described in Fig. 1d. 1x indicates normal size embryo, resulting from aggregating 8 cells, 2x indicates double-sized embryo, resulting from aggregating 2x intact 8-cell stage embryos. **(H)** Box plots showing the relative contribution of GFP+ (green, wt) and GFP-, *Gata6^−/−^* cells (magenta) to the embryo for each of the initial % of GFP+ cells, as shown in (G) Dark green spots represent double embryos (2x). **(I)** Box plots showing the relative contribution of GFP+ (green, wt) and GFP-, *Gata6^−/−^* cells (magenta) to the ICM for each of the initial % of GFP+ cells, as shown in (G) **(J)** Optical cross sections through embryos shown in (G) and immunostained for the markers indicated. **(K)** Box plots showing the relative contribution of GFP+ (green, wt) and GFP-, *Gata6^−/−^* (magenta) cells to each the PrE and the epiblast for embryos grouped by the final fraction of GFP+ ICM cells, as in (E) and as indicated on the x-axis. **(L)** Stacked bar plots showing the ICM composition in *Gata6^−/−^* ↔ GFP+ chimeras, grouped by the final fraction of GFP+ ICM cells, as in (K) and as indicated on the x-axis. Color coding is indicated. **(M)** Box plots showing the % of GFP+ or GFP-cells (both wild type) that adopt EPI/DN identity in wt ↔ GFP+ chimeras, grouped by the final fraction of GFP+ ICM cells, as in (E,K) and as indicated on the x-axis. **(N)** Box plots showing the % of GFP+ (wt) or GFP-, *Gata6^−/−^* cells that adopt EPI/DN identity in *Gata6^−/−^* ↔ GFP+ chimeras, grouped by the final fraction of GFP+ ICM cells, as in (M). In all box plots whiskers span 1.5x the inter quartile range (IQR) and open circles represent outliers (values beyond 1.5x IQR). Cross indicates the arithmetic mean and each dot represents one embryo. All embryos labeled for NANOG (red), GATA6 (blue) and either CDX2 (controls) or GFP (chimeras) (green). All optical cross sections are 5µm maximum intensity projections. Total cell counts are indicated for each embryo within the merged images. PrE: Primitive Endoderm, DP: Double Positive (for NANOG and GATA6), EPI: Epiblast, EPI-lo: low NANOG epiblast, DN: Double Negative (for NANOG and GATA6). Scale bars = 20µm.

**Supplementary Figure 3 (related to Figure 2).**
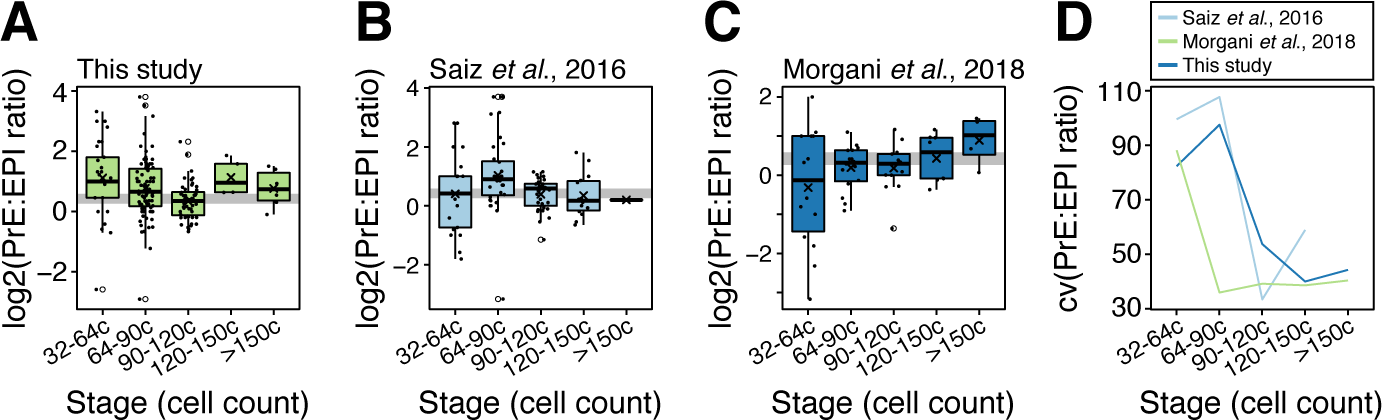
**(A-C)** Box plots showing the ratio of PrE to epiblast cells in embryos at sequential stages of blastocyst development for embryos collected in this study (A), thos analyzed in (Saiz et al., 2016b) (B) and those analyzed in (Morgani et al., 2018b) (C) (shown also in Fig. S1A-B). In all box plots whiskers span 1.5x the inter quartile range (IQR) and open circles represent outliers (values beyond 1.5x IQR). Crosses indicate the arithmetic mean and each dot represents one embryo. **(D)** Coefficient of variation of the PrE:epiblast ratio over developmental time in the datasets shown in (A), (B) and (C), as indicated.

**Supplementary Figure 4 (related to Figure 3).**
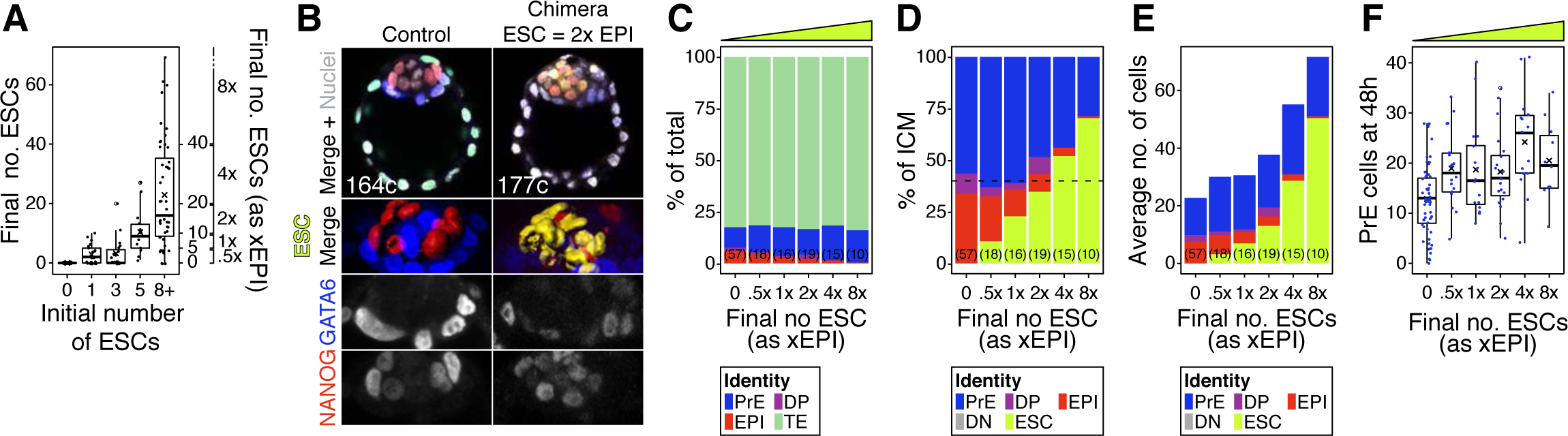
**(A)** Box plots indicating ESC contribution to chimeras. Number of ESCs aggregated with morulae is shown against the final number of ESC in the chimeric embryo after 48h in culture. Right y axis shows epiblast-equivalent size bins used to categorize chimeric embryos (xEPI). **(B)** Optical cross sections through representative immunofluorescence images of a control and chimeras with 2xEPI- and 4xEPI-equivalent ESC compartments. Embryos are labelled for NANOG (red) and GATA6 (blue) to identify all ICM cell types. ESCs are shown in yellow. Lower panels show magnifications of the ICM, with all markers overlaid, for each individual marker as grayscale and for ESCs as 3D surface renders. Total cell count for each embryo is shown within the merged panel. **(C)** Stacked bar plot showing the overall lineage distribution of host cells in each group of embryos shown in Fig. 3, grouped by the final size of the ESC compartment (also represented by the yellow wedge) **(D)** Stacked bar plot showing the average relative ICM composition of each group of chimeras shown in (C). **(E)** Stacked bar plot showing the average number of cells in each ICM population for each group of embryos shown in (D). **(F)** Box plot showing the size of the PrE at the end of the experiment in each group of embryos. Yellow wedge represents the increasing amount of ESCs in each group. Color coding is indicated. Number of embryos in each group is indicated in parenthesis. In all box plots whiskers span 1.5x the inter quartile range (IQR) and open circles represent outliers (values beyond 1.5x IQR). Cross indicates the arithmetic mean and each dot represents one embryo. All optical cross sections are 5µm maximum intensity projections. PrE: Primitive Endoderm, DP: Double Positive (for NANOG and GATA6), EPI: Epiblast, ESC: embryonic stem cell. Scale bars = 20µm.

**Supplementary Figure 5 (related to Figure 5).**
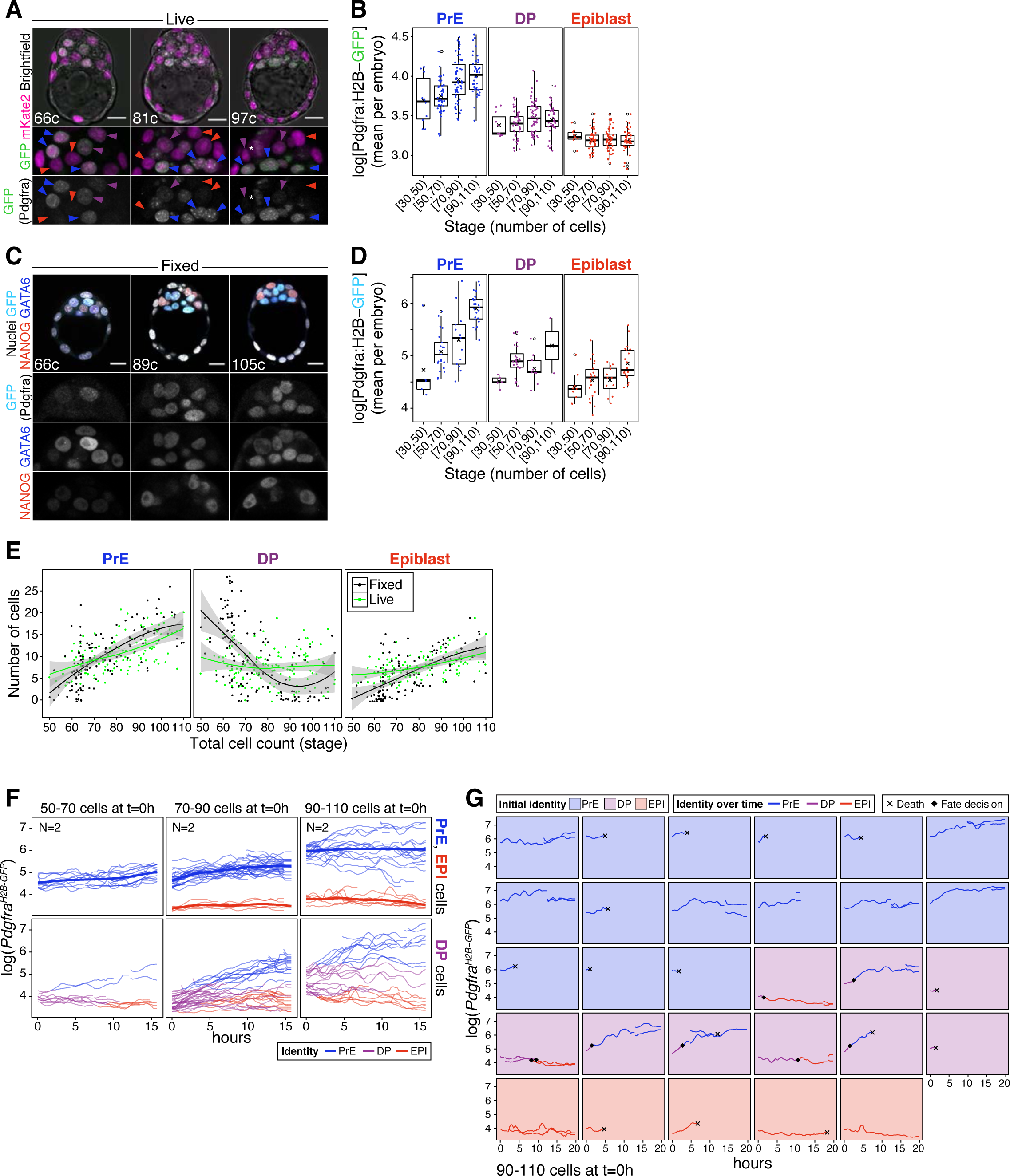
**(A)** Optical cross sections through representative live blastocysts at sequential developmental stages (total cell count is indicated within each image). Lower panels show magnifications of the ICM for both markers, as indicated, and for GFP alone in grayscale. Colored arrowheads point at nuclei considered to be PrE (blue arrowheads), DP (purple arrowheads) or epiblast (red arrowheads) based on GFP level. **(B)** Box plots showing the average level of GFP (*Pdgfra*) per embryo, in each ICM cell type, for each developmental stage considered. **(C)** Optical cross sections through representative immunofluorescence images of reference *Pdgfra^H2B-GFP/+^* littermates fixed upon collection and labelled for NANOG (red) and GATA6 (blue) to identify ICM cell types. Lower panels show ICM magnifications for each marker in grayscale, as indicated. **(D)** Box plots showing the average level of GFP (*Pdgfra*) per embryo, in each ICM cell type (as determined by NANOG and GATA6 expression), for each developmental stage considered – for in embryos like those in (C). **(E)** Growth curves for each ICM population over time. Black lines and dots show numbers corresponding to fixed samples, where cell identities were assigned automatically based on relative NANOG and GATA6 levels, as described in the Methods. Green lines and dots show numbers corresponding to live samples, where cell identities were assigned manually based on GFP (*Pdgfra*) levels alone, as described in the Methods. Curves are local regression lines for each subset of data, fitted using the LOESS method. **(F)** Temporal dynamics of cell fate specification in live embryos. Expression levels of the PrE reporter allele *Pdgfra^H2B-GFP^* (Hamilton et al., 2003) are shown over time for individual ICM cells in intact embryos at sequential stages of development. Reporter expression was assessed using time lapse imaging over a 15h time window. Top panels show dynamics of reporter expression in cells classified as PrE or epiblast at the beginning of the movie. Smoothing curves for each PrE and epiblast are shown as thicker lines and color coded. Bottom panels show dynamics of reporter expression in cells classified as DP (progenitors) at the beginning of the movie. DP cells become PrE or epiblast over the course of the movie, as determined by *Pdgfra^H2B-GFP^* expression. **(G)** *Pdgfra* expression dynamics for individual cells in one embryo from (F) imaged from the 90-110 cell stage onwards. Panels are color coded for the initial identity of the cell. Traces are color coded for identity over time. Acquisition of PrE or epiblast identity is denoted by a black diamond. Black cross indicates cell death. In all box plots whiskers span 1.5x the inter quartile range (IQR) and open circles represent outliers (values beyond 1.5x IQR). Cross indicates the arithmetic mean and each dot represents one embryo. PrE: Primitive Endoderm, DP: Double Positive (for NANOG and GATA6), EPI: Epiblast. Scale bars = 20µm.

**Supplementary Figure 6 (related to Figure 6).**
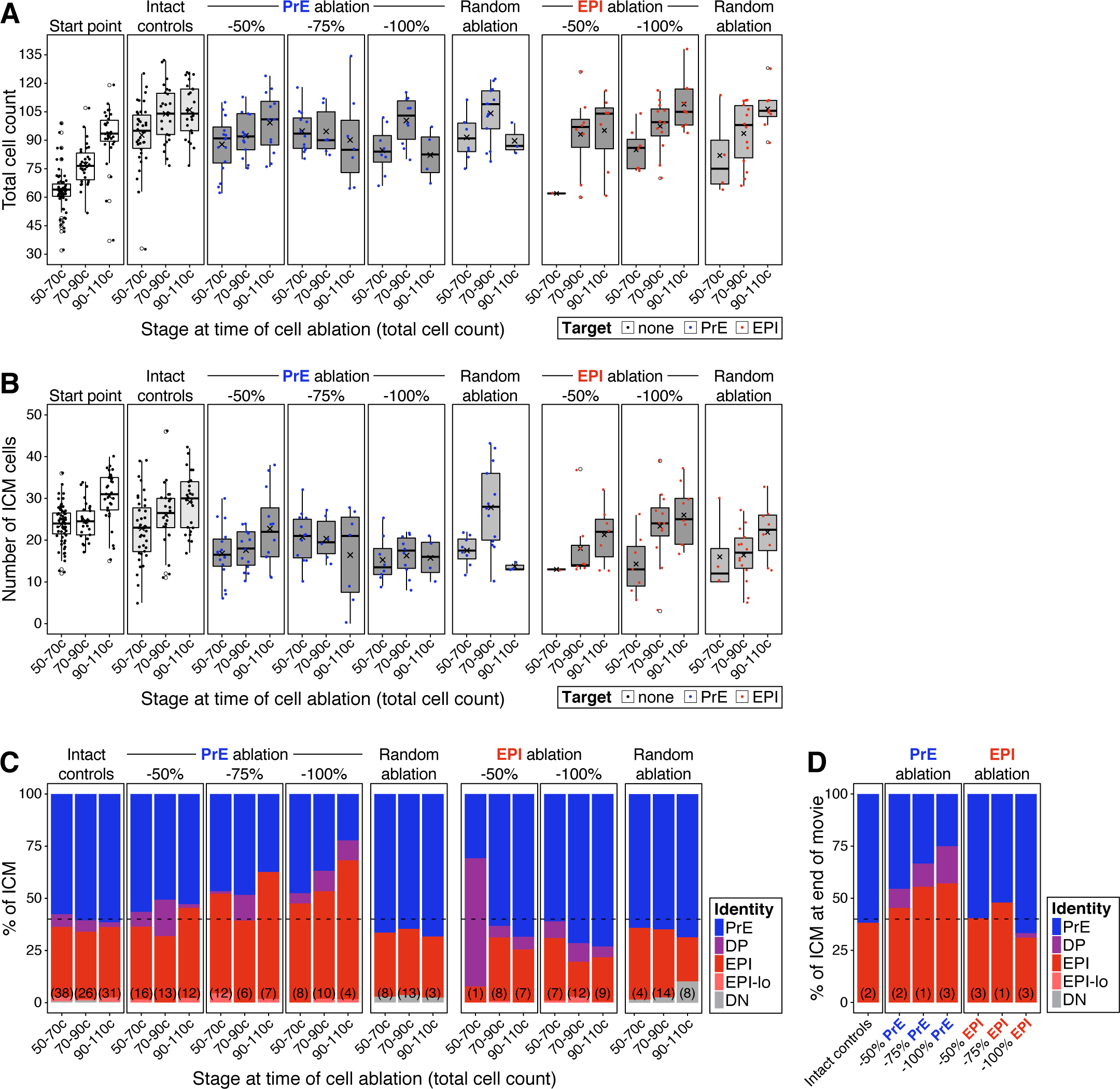
**(A)** Box plots showing the total cell count of embryos in each of the experimental groups shown in Fig. 6, at the end of the experiment. Embryos are binned by developmental stage at the start of the experiment, as shown on the x-axis. Start point comprises reference littermates fixed at the beginning of the experiment. Intact controls are embryos in which no cell was targeted. Random controls are embryos in which randomly chosen ICM cells were targeted, irrespective of their identity, in equivalent numbers to the −100% group for each lineage targeted (see Methods). Embryos in which the PrE or epiblast were targeted are split by the fraction of the lineage eliminated (50-100%, as indicated). **(B)** Box plots showing the total number of ICM cells in each of the experimental groups shown in (A). **(C)** Stacked bar plots showing the relative ICM composition in each of the experimental groups shown in Fig. 6c-d and in (A-B) above. The number of embryos in each group is indicated in parenthesis. Cell types are color-coded as indicated. **(D)** Stacked bar plots showing the relative ICM composition at the end of the movie in embryos shown in Fig. 6e, for each of the treatments indicated on the x-axis. Data in (D) corresponds to embryos in which ablation was performed at the 70-90 cell stage and in which all or most of the ICM cells could be tracked throughout the movie. Cell types are color-coded as indicated. In all box plots whiskers span 1.5x the inter quartile range (IQR) and open circles represent outliers (values beyond 1.5x IQR). Cross indicates the arithmetic mean and each dot represents one embryo. PrE: Primitive Endoderm, DP: Double Positive (for NANOG and GATA6), EPI: Epiblast.

**Supplementary Figure 7 (related to Figure 6).**
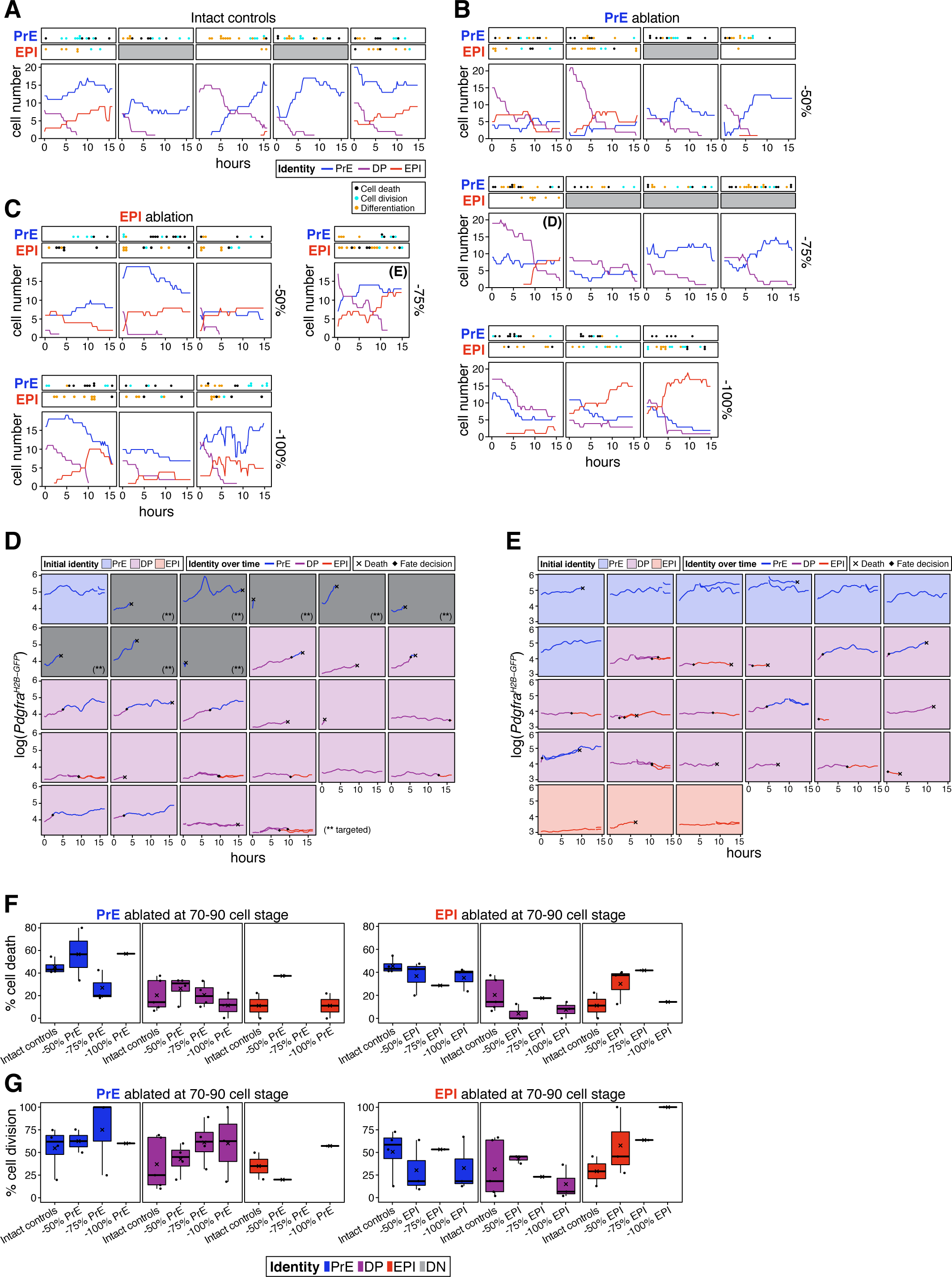
**(A)** Growth curves for each of the ICM populations in each of the intact embryos shown in Fig. 6E. Cell types are color-coded. Top panels indicate cellular events in the PrE or epiblast populations: black dots represent cell death, cyan dots represent cell divisions, and orange dots represent a progenitor cell adopting either PrE or epiblast identity, respectively. Gray panels indicate no data is available **(B)** Growth curves for each of the ICM populations in each of the embryos shown in Fig. 6E in which the PrE was targeted, in the fractions indicated. **(C)** Growth curves for each of the ICM populations in each of the embryos in which the epiblast was targeted, shown in Fig. 6E. All embryos in (A-C) were manipulated at the 70-90 cell stage and live imaged for the first 15h of the 24h culture after ablation. **(D)** Temporal changes in *Pdgfra* expression and cell identity for all ICM cells tracked in an embryo in which 75% of the PrE was eliminated when it had 70-90 cells – labelled as “(D)” in (B). **(E)** Temporal changes in *Pdgfra* expression and cell identity for all ICM cells tracked in an embryo in which 75% of the epiblast was eliminated when it had 70-90 cells – labelled as “(E)” in (C). Each panel displays one cell. Panels are color coded for the initial identity of the cell, as indicated. Traces are color coded for identity over time. Acquisition of PrE or epiblast identity is denoted by a black diamond. Black cross indicates cell death. **(F)** Box plots showing frequency of cell death among intact cells of each ICM cell type, as color-coded. Each box represents one experimental group (Intact controls, or embryos in which increasing fractions of the corresponding lineage were eliminated, as indicated on the x-axis). Left plots correspond to PrE ablation, right plots to epiblast ablation, as indicated. **(G)** Box plots showing the frequency of cell division among intact cells of each ICM cell type, as color coded. Each box represents one experimental group (Intact controls, or embryos in which increasing fractions of the corresponding lineage were eliminated, as indicated on the x-axis). Left plots correspond to PrE ablation, right plots to epiblast ablation, as indicated. Color coding is indicated. In all box plots whiskers span 1.5x the inter quartile range (IQR) and open circles represent outliers (values beyond 1.5x IQR). Cross indicates the arithmetic mean and each dot represents one embryo. PrE: Primitive Endoderm, DP: Double Positive (for NANOG and GATA6), EPI: Epiblast.

**Supplementary Figure 8 (related to Figure 7).**
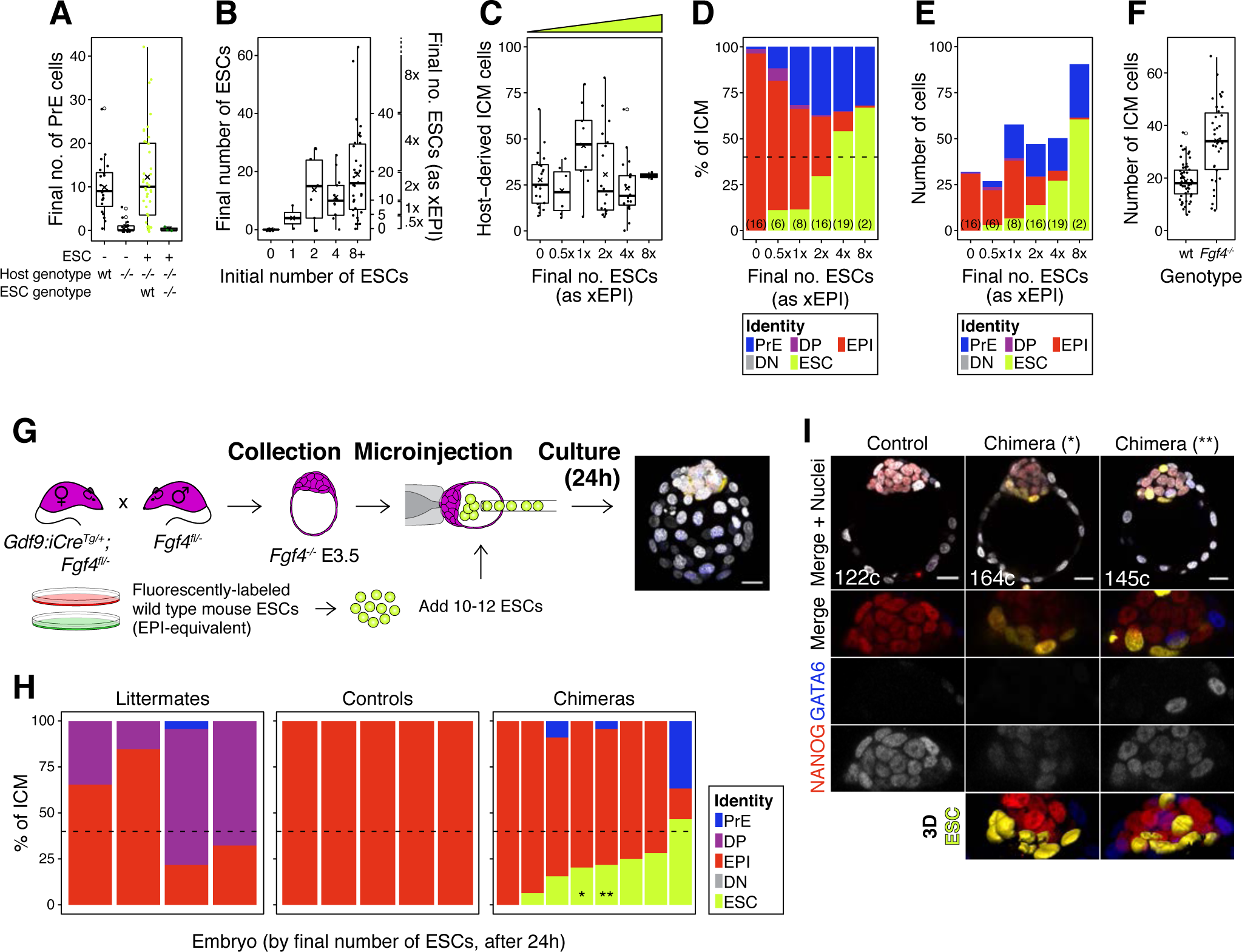
**(A)** Box plot indicating the number of PrE cells in wild type control embryos, *Fgf4^−/−^* controls, *Fgf4^−/−^* chimeras carrying wild type ESCs and *Fgf4^−/−^* chimeras carrying GFP-tagged, *Fgf4^−/−^* ESCs, as indicated. **(B)** Box plots indicating ESC contribution to chimeras. Number of ESCs aggregated with morulae is shown against the final number of ESC in the chimeric embryo after 48h in culture. Right y-axis shows epiblast-equivalent size bins used to categorize chimeric embryos. **(C)** Box pot showing the size of the host-derived ICM component at the end of the experiment in each group of embryos. **(D)** Stacked bar plots showing the average relative ICM composition of *Fgf4^−/−^* embryos carrying wild type ESCs (shown in Fig. 7D), binned by the final size of the ESC compartment. **(E)** Stacked bar plots showing the average number of each ICM cell type for each group of chimeras shown in (D) and Fig. 7D. **(F)** Box plot showing the ICM size of wild type control embryos and *Fgf4^−/−^* control embryos. **(G)** Experimental design for blastocyst injection. Embryos were recovered from crosses equivalent to those in Fig. 7A, at the mid-blastocyst stage (∼60-80 cells) and 10-12 ESCs injected into the blastocyst cavity before allowing the embryos to develop for 24-30h in culture, until a stage equivalent to ∼E4.5 days post-fertilization. **(H)** Stacked bar plots showing ICM composition for individual embryos like those shown in (I), as indicated. Each bar represents the ICM of one embryo and bars are arranged by absolute number of ESCs present. Stars (*, **) denote the bars corresponding to the chimeras shown in (I). Color coding is indicated. **(I)** Optical cross sections through representative chimeras and control embryos cultured from the blastocyst stage and labelled for NANOG and GATA6 to identify all ICM cell types. The progeny of the introduced ESCs is labelled in yellow. These cells failed to mix with host ICM cells and tended to remain at the ICM surface, suggesting that differences in adhesion between cell types are necessary for cell mixing (see also Movies S10-11). Lower panels show magnifications of the ICM, with all markers overlaid, for each individual marker in grayscale and for ESCs as 3D surface renders. Total cell count is indicated for each embryo. All optical cross sections are 5µm maximum intensity projections. In all box plots whiskers span 1.5x the inter quartile range (IQR) and open circles represent outliers (values beyond 1.5x IQR). Cross indicates the arithmetic mean and each dot represents one embryo. Yellow wedges represent the increasing amount of ESCs in each group. PrE: Primitive Endoderm, DP: Double Positive (for NANOG and GATA6), EPI: Epiblast, DN: Double Negative (for NANOG and GATA6), ESC: embryonic stem cell. Scale bars = 20µm.

